# Novel COX-2–Targeted Nanobodies for Molecular Endoscopic Imaging of Colorectal Adenomas

**DOI:** 10.64898/2026.05.16.724741

**Authors:** Md. Jashim Uddin, Shu Xu, Michael C. Goodman, Ansari M. Aleem, Hiroaki Niitsu, Kristie L. Rose, Brenda C. Crews, Surajit Banerjee, Carlisle R. DeJulius, Ella N. Hoogenboezem, Philip J. Kingsley, Michelle L. Reyzer, Jamie Klendworth, Matthew Milad, Shuyang Lin, Brian E. Wadzinski, Benjamin W. Spiller, Craig L. Duvall, Robert J. Coffey, Lawrence J. Marnett

**Author notes:** **Corresponding Authors** Md. Jashim Uddin, **E-mail:**, and Lawrence, J. Marnett, **E-mail:**.

## Abstract

Colorectal cancer (CRC) is one of the leading causes of cancer-related mortality in men and women. Timely detection and diagnosis are key to management of CRC, which is under-diagnosed because colorectal aberrant crypt foci, hyperplastic polyps, and microadenomas are often missed with conventional colonoscopy. The enzyme cyclooxygenase-2 (COX-2) is overexpressed in early stages of colorectal carcinogenesis and plays an important regulatory role in the process, suggesting that it could be a valuable target for enhanced imaging of nascent disease. Thus, we have generated an alpaca-derived library of 73 COX-2-specific nanobody clones. Here, we describe one such nanobody, F9-K45Q-K77Q-ROX, in which two native lysine residues have been mutated followed by conjugation to a fluorophore at the *N*-terminus with retention of COX-2-selective binding. The site of fluorophore conjugation and COX-2 binding affinity of F9-K45Q-K77Q-ROX were determined by proteomic and microscale thermophoretic analyses, respectively. In cell culture studies using 1483 human head and neck squamous cell carcinoma cells, F9-K45Q-K77Q-ROX accumulated inside cells and bound to intracellular COX-2, as visualized by fluorescence microscopy. *In vivo* pharmacokinetic, and toxicological analyses revealed that F9-K45Q-K77Q-ROX is detectable in circulation with a plasma half-life of 17.9 min and there is no short-term toxicity associated with single injections of 10 mg/kg, 20 mg/kg, or 40 mg/kg doses at 24 h post-administration. Noninvasive *in vivo* fluorescence endoscopic imaging validated tumor-specific accumulation of F9-K45Q-K77Q-ROX in azoxymethane/dextran sodium sulfate-induced colorectal adenomas in mice. This work demonstrates the first COX-2-targeted nanobodies including a fluorescent derivative that offers significant promise for targeted endoscopic imaging of COX-2-expressing neoplasms.

**Significance Statement:** Current colorectal cancer screening procedures, such as white-light colonoscopy, chromoendoscopy, and narrow-band imaging aim to detect solid colon tumors and precursor lesions. However, these methods tend to detect only raised solid tumors and mature cancers, whereas precursor lesions, such as aberrant crypt foci, hyperplastic polyps, and small adenomas are frequently missed. To address the need for better visualization of early lesions, we developed a library of alpaca-derived nanobodies targeted to cyclooxygenase-2 (COX-2), an enzyme that is overexpressed in colorectal adenomas. COX-2-targeted nanobodies bearing a fluorescent tag accumulate and are retained in colonic adenomas, facilitating their endoscopic visualization. This novel COX-2-targeted nanobody platform may also be valuable for early detection of other neoplastic diseases in which COX-2 overexpression occurs. (Word counts 119, limit 120)

## Introduction

Technologies for imaging colorectal cancer (CRC) in animal models and patients have been significantly improved through advances in instrumentation, enabling high-resolution visualization of tissue structure (1-3). Furthermore, decades of basic and clinical research have identified several molecular biomarkers that may be valuable for assessing susceptibility to CRC, and CRC progression, as well as dissecting molecular mechanisms of CRC pathophysiology in preclinical studies (4-14). Early detection and diagnosis is key to effective management of CRC. Current CRC imaging strategies, including white light colonoscopy, chromoendoscopy, and narrow-band imaging aim to detect and enable surgical resection of solid tumors and precursor lesions (15-18). Using these methods, raised solid tumors and mature CRC can typically be visualized and removed, whereas aberrant crypt foci and hyperplastic polyps are frequently missed (19-24). This miss rate is clearly an unmet medical need that warrants critical development of novel strategies enabling early diagnostic imaging of CRC in preclinical and clinical settings.

COX-2 overexpression is one of the major driving forces of colorectal tumorigenesis (25-29), suggesting that COX-2 is a promising imaging target for the development of diagnostic agents for early CRC. We and others have shown *in vivo* imaging of COX-2 in multiple animal models of premalignant and malignant tumors and inflammation using small molecule fluorescently tagged or radiolabeled agents (30-40). These studies provide proof-of-principle for targeted molecular imaging of COX-2 in *in vivo* pathogenesis.

All of the COX-2-targeted imaging agents to date are small molecules that are based on COX-2-selective inhibitors that bind in the cyclooxygenase active site. Fluorophores are tethered to the inhibitor core or radioisotopes (^18^F or ^123^I) are incorporated into the inhibitor framework. The incorporation of the fluorescent or radiochemical functionality must be consistent with the structure-activity for the particular inhibitor moiety against COX-2. Also, since COX-2 is an intracellular protein, the imaging agent must possess sufficient hydrophobicity to cross cellular membranes. These restrictions limit the functional groups and modalities that can be used for COX-2-targeted imaging.

To date, no imaging agents have been developed that bind to the surface of the COX-2 protein outside of the cyclooxygenase active site. COX-2-specific monoclonal antibodies exist, but their size (150 kDa) may limit their ability to penetrate tumor tissue or to cross cellular membranes. Nanobodies (∼15 kDa) are the smallest functional antigen-binding fragments and are derived from naturally occurring heavy-chain-only antibodies that were first identified in the blood of llamas (41-43). Their size allows increased tissue penetration compared to that of monoclonal antibodies (42), and they are water-soluble (44), unlike many small molecule probes. These factors suggest that nanobodies would be well-suited to formulation for systemic dosing (45, 46) although their ability to cross cellular membranes is uncertain. Herein, we report the discovery of a COX-2-specific nanobody and a nanobody-based fluorescence imaging agent capable of targeting COX-2 in intact cells and mouse colorectal adenomas *in vivo*. The novel immunoprobe combines structural and functional imaging readouts obtained with fluorescence-aided colonoscopy that provide powerful diagnostic information for early detection and improved management of colorectal carcinogenesis. (Word counts: 496)

## Results

A flowchart (47, 48) for generating and identifying anti-COX-2 nanobodies is shown in Fig. 1A. Peripheral blood mononuclear cells were purified from alpacas immunized with murine COX-2 (mCOX-2), and mRNA was isolated from these cells. A library of single chain variable regions was amplified by reverse transcription polymerase chain reaction and cloned into a phagemid vector that resulted in fusion of the antibody gene to a coat protein of the m13 bacteriophage for expression in *E. coli*. Screening of the library revealed a number of promising clones. One of these – designated “F9” – was isolated, expressed in *E. coli*, and purified.

**Figure 1.**
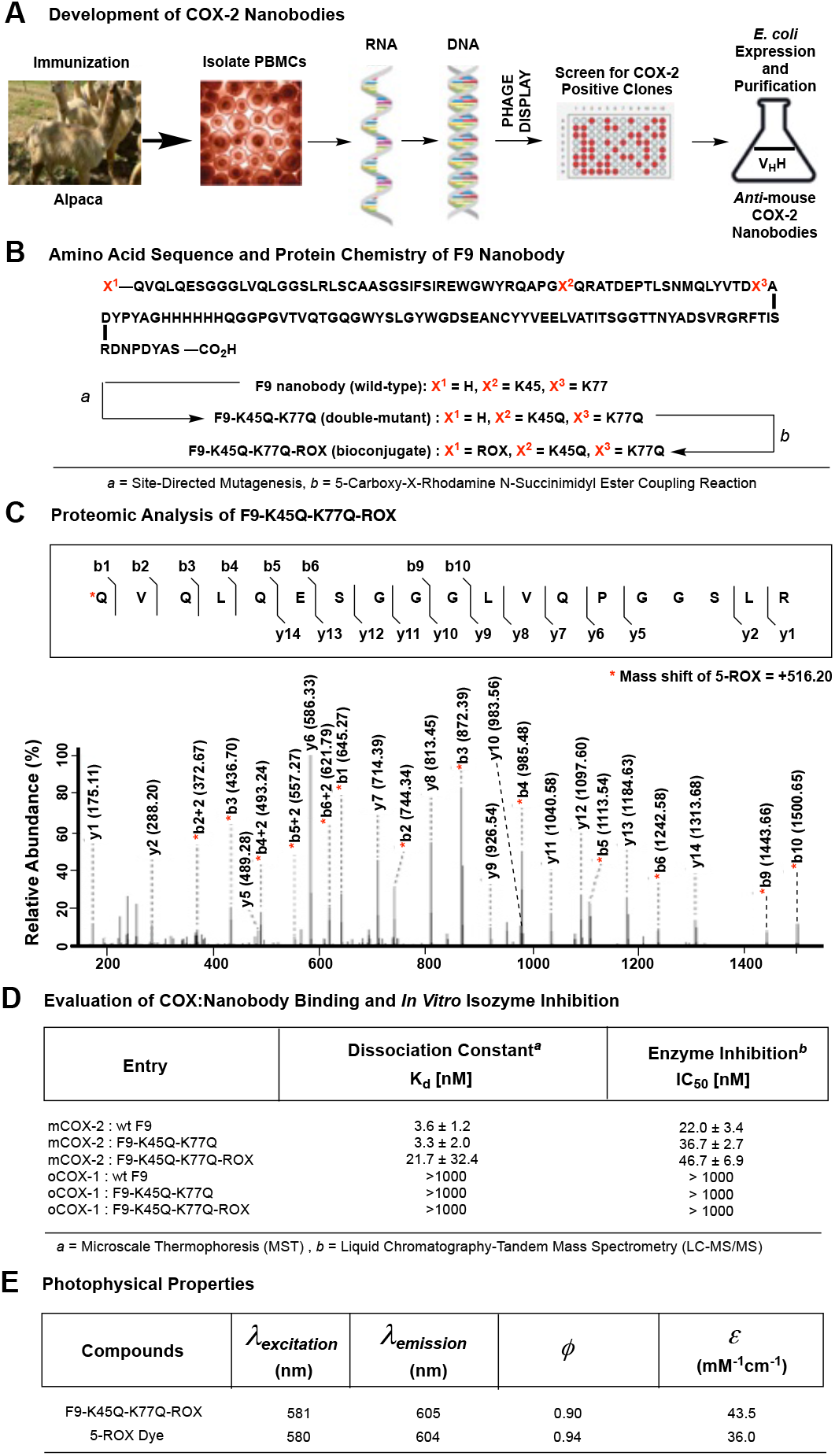
(A) Schematic representation of the development of anti-mouse COX-2 nanobodies; (B) transformation of F9 into F9-K45Q-K77Q double mutant by site-directed mutagenesis followed by the conjugation of 5-ROX-NSE to give F9-K45Q-K77Q-ROX; (C) proteomic analysis of F9-K45Q-K77Q-ROX by in-gel trypsin digestion followed by tandem LC-MS/MS spectroscopy of tryptic peptide residues 1-18 labeled with 5-ROX (mass shift +516.20) at the N-terminus, where the observed b- and y-type product ions are assigned to their corresponding m/z peaks in the mass spectrum. The amino acid sequence is provided above the annotated spectrum, and sites of amide bond fragmentation are indicated with inter-residue brackets. Asterisks indicate the b-type product ions shifted in mass due to the addition of 5-ROX; (D) in vitro biochemical evaluation of COX binding affinity of F9, F9-K45Q-K77Q or F9-K45Q-K77Q-ROX nanobodies by microscale thermophoresis and COX inhibition by F9, F9-K45Q-K77Q or F9-K45Q-K77Q-ROX nanobodies; (E) photophysical properties of F9-K45Q-K77Q-ROX compared to free 5-ROX dye.

The sequence of F9 is displayed in Fig. 1B. It contains two lysine residues at the 45^th^ and 77^th^ positions and a glutamine residue at the *N*-terminus. Site-specific fluorophore conjugation at the free primary amino-group of the *N*-terminal residue required mutation of the two lysine residues to amino acids lacking a free amino group. Thus, we used standard site-directed mutagenesis methodology (32, 49, 50) to generate the F9-K45Q-K77Q double mutant, which was expressed in *E. coli* and purified. Then, 5-carboxy-X-rhodamine (5-ROX) was conjugated by reacting F9-K45Q-K77Q with 5-ROX *N*-succinimidyl ester (Fig. 1B). An analytical HPLC confirmed the complete consumption of F9-K45Q-K77Q in the reaction mixture (Fig. S1). A size-exclusion column (SEC) chromatography of the reaction mixture on a UPLC system afforded the fluorescent F9-K45Q-K77Q-ROX nanobody with a degree of labeling of >99% (Fig. S7) determined by a NanoDrop 3300 Fluorospectrometer recording fluorescence measurements in a typical protein chemistry preparation.

The site of 5-ROX conjugation on the F9-K45Q-K77Q-ROX nanobody was determined by proteomic analysis following in-gel trypsin digestion. The fragmentation pattern of the nanobody (Fig. 1C) revealed that 5-ROX conjugation had occurred at the *N*-terminus.

The F9 nanobody was generated using apo-COX-2, so the screening assays used to identify the most active clones also employed the apo-enzyme. We determined the dissociation constants for the binding of Wt-F9, F9-K45Q-K77Q, and F9-K45Q-K77Q-ROX nanobodies to apo mCOX-2 by a microscale thermophoresis (MST) assay (51). The assay identified Wt-F9, F9-K45Q-K77Q and F9-K45Q-K77Q-ROX as high affinity COX-2-binding nanobodies with dissociation constants (K_d_) of 3.6 ± 1.2 nM, 3.3 ± 2.0 nM and 21.7 ± 32.4 nM, respectively (Figs. 1D and S2). When the binding affinity to oCOX-1 was evaluated, the nanobodies exhibited dissociation constants of > 1 μM, suggesting a high degree of COX-2-selectivity.

Using a photon counting fluorometer, we determined the photophysical properties of F9-K45Q-K77Q-ROX, using 5-ROX as a well-characterized standard fluorescent compound for comparison. In this assay, F9-K45Q-K77Q-ROX exhibited quantum yield (Φ_fl_ = 0.9 at 605 nm) and extinction coefficient (ε = 43.5 mM^-1^cm^-1^ at 581 nm) values that were comparable to the values obtained for 5-ROX (Φ_fl_ = 0.94 at 604 nm, ε = 36 mM^-1^cm^-1^ at 580 nm) (Fig. 1E).

To elucidate the molecular basis of binding, we complexed and co-crystallized the F9 nanobody with purified mCOX-2. The structure of the F9:apoCOX-2 crystal complex was solved by X-ray crystallography at 2.15 Å resolution (Fig. 2A). COX-2 is a homodimer of subunits that each contain a cyclooxygenase active site and a peroxidase active site separated by a heme prosthetic group. The cyclooxygenase active site catalyzes the conversion of arachidonic acid (AA) to the hydroperoxyendoperoxide intermediate, prostaglandin G_2_ (PGG_2_). The hydroperoxide of PGG_2_ is then reduced at the peroxidase active site, yielding prostaglandin H_2_ (PGH_2_), which subsequently serves as substrate for other enzymes in the PG synthetic pathway. The crystal structure revealed that one molecule of F9 binds at the peroxidase active site of each monomer of the COX-2 protein. The F9 nanobody binds to COX-2 via several key hydrophobic interactions and numerous direct and solvent-mediated hydrophilic interactions. Residues of F9, including L107, W104, Y105, D102, S32, F31, S55, T52, N60, T54, and T58 interact through 17 bridging water molecules with H207, Q289, R222, N382, H214, E457, S174, and R456 of COX-2 (Fig. 2B). It should be noted that an unusual bridging structure can be found in which D102 of F9 and N382 of COX-2 interact through 4 water molecules. The structure provides the molecular basis for the observed affinity of F9 for mCOX-2.

**Figure 2.**
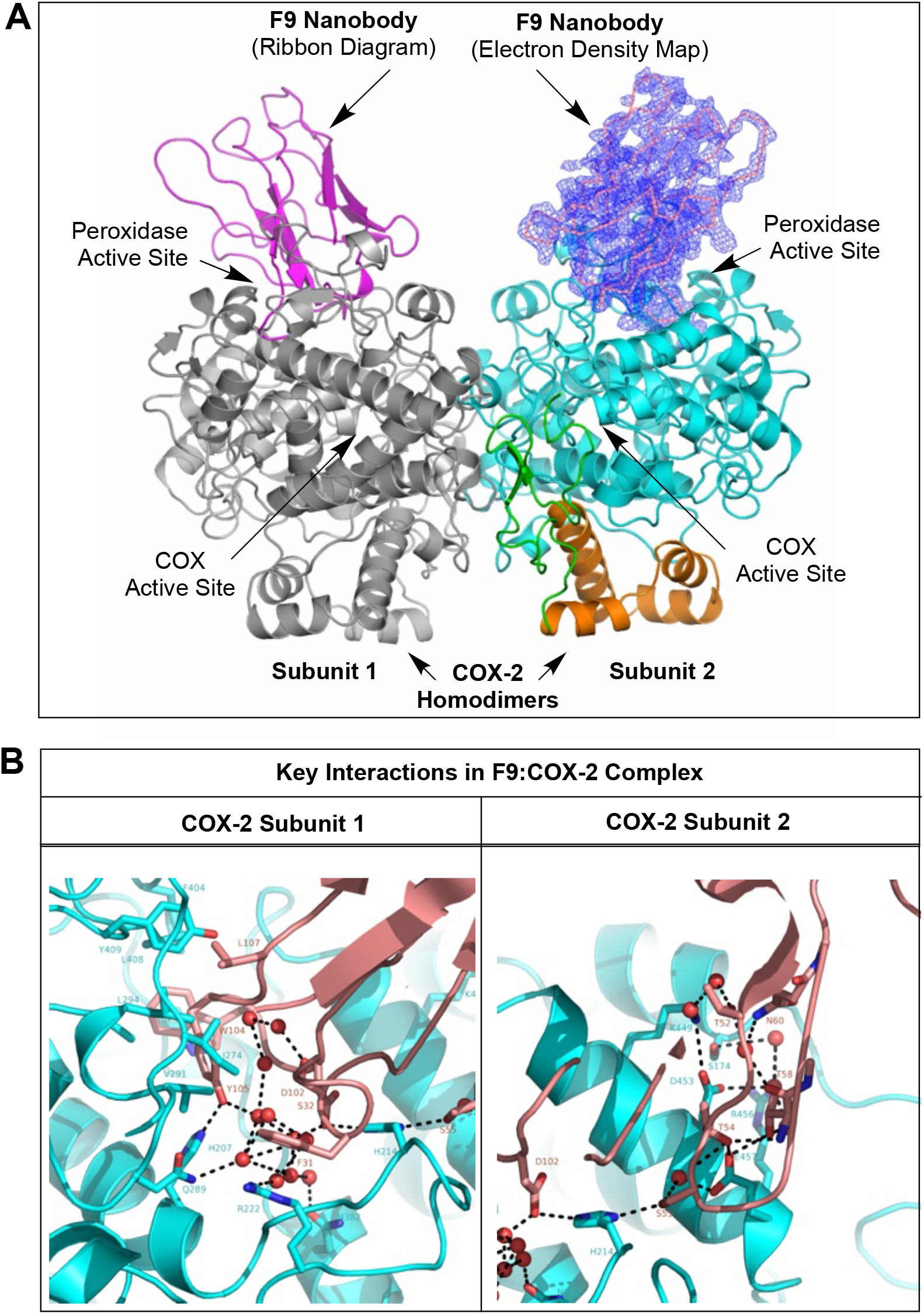
(A) X-Ray co-crystal structure of the F9:COX-2 complex solved at 2.15 Å resolution (PDB ID: 8ET0), revealing that the F9 nanobody binds at the peroxidase active site of each subunit of COX-2 homodimer; (B) key interactions between F9 and the COX-2 homodimer within the F9:COX-2 co-crystal complex.

The F9:COX-2 complex with the structure of holo-COX-2 *(PDB ID: 8ET0)* indicates that the nanobody partially fills and obstructs the heme binding site (Figs. 2A,B & Table S1). This finding is consistent with our failure to generate crystals of the holoenzyme in complex with the nanobody. Thus, it appears that heme and the nanobody cannot bind simultaneously to the same COX-2 subunit. Although crystal structures of holo-COX-2 reveal heme in both subunits, data suggest that the enzyme in solution binds heme with high affinity in only one subunit (52). Thus, the enzyme could conceivably have heme bound to one subunit while F9 is simultaneously bound to the other.

To further explore the possible interactions between F9 and heme binding, we assessed the effects of the F9, F9-K45Q-K77Q and F9-K45Q-K77Q-ROX nanobodies on the conversion of AA to PGE_2_ and PGD_2_ (the nonenzymatic rearrangement products of PGH_2_) by purified recombinant mCOX2 in a standard enzyme activity assay. We found that when apoCOX-2 was first reconstituted with heme prior to incubation with the nanobody followed by addition of substrate, the nanobodies had no effect on enzyme activity. In contrast, if we first incubated apoCOX-2 with varying concentrations of test nanobodies followed by the addition of heme and then AA, production of PGE_2_ and PGD_2_ was inhibited. The F9, F9-K45Q-K77Q and F9-K45Q-K77Q-ROX nanobodies exhibited IC_50_ values of 22.0 ± 3.4 nM, 36.7 ± 2.7 nM, and 46.7 ± 6.9 nM, respectively (Fig. S3). The results suggest that the nanobody blocks the binding of heme to the apoenzyme, but heme already bound to the enzyme is not readily displaced by the nanobody.

We evaluated F9-K45Q-K77Q-ROX’s ability to detect COX-2 in human head and neck squamous cell carcinoma (1483 HNSCC) cells. A human ovarian cancer (OVCAR-3) cell line that does not express COX-2 but overexpresses the alternative COX isoform, COX-1, served as a negative control. Cell lines were treated with F9-K45Q-K77Q-ROX and washed with serum-containing medium followed by fluorescence microscopy. As revealed by optical imaging, 1483 HNSCC cells were reproducibly labeled by F9-K45Q-K77Q-ROX (Fig. 3A), whereas OVCAR-3 cells exhibited negligible fluorescence (Fig. 3B). Cell fluorescence was quantified on a BioTek Synergy Mx Plate Reader showing a statistically significant fluorescence difference between COX-2-expressing and COX-1-expressing cell lines (Fig. 3C). These observations support the hypothesis that F9-K45Q-K77Q-ROX is COX-2-specific in cells.

**Figure 3.**
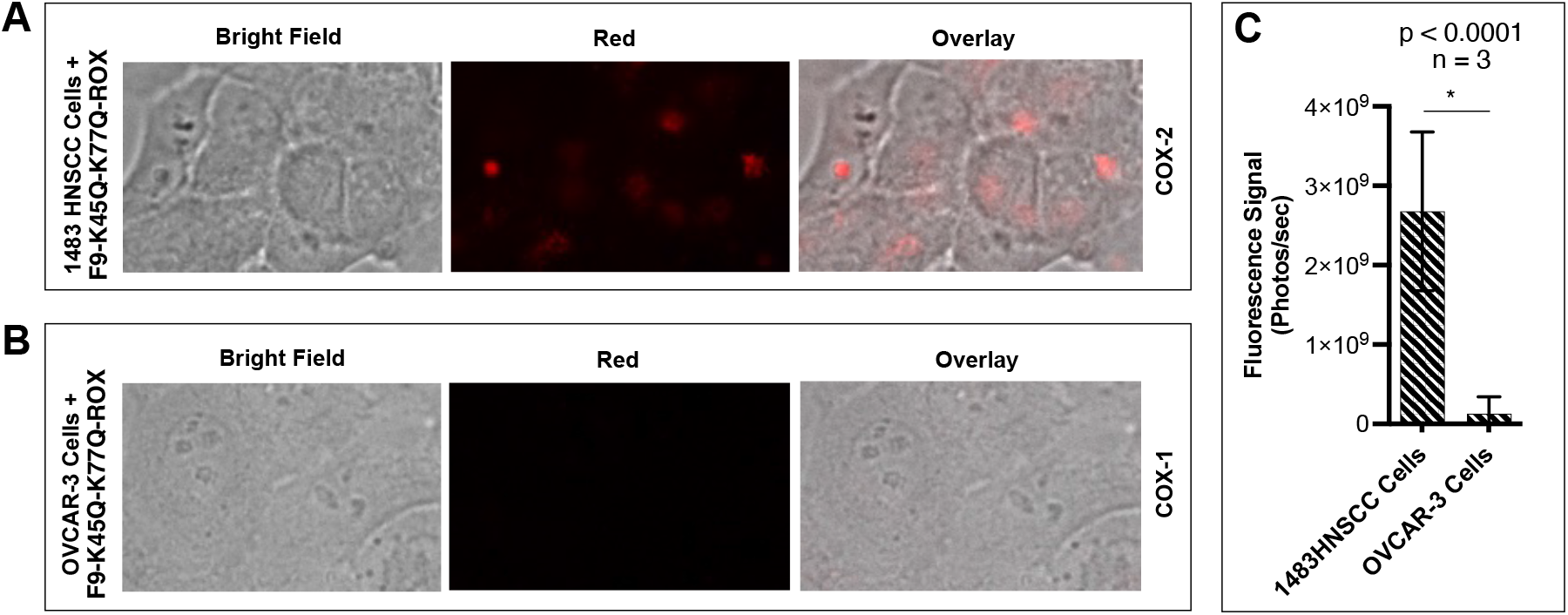
Fluorescence microscopy of 1483 HNSCC cells (COX-2 expressing) (A) or (B) OVCAR-3 cells (COX-1 expressing) (B) treated with F9-K45Q-K77Q-ROX, (C) measurement of fluorescence signals in 1483 HNSCC cells versus OVCAR-s cells treated with the F9-K45Q-K77Q-ROX nanobody.

We determined the *in vivo* plasma half-life of F9-K45Q-K77Q-ROX in CD1 mice using 5-ROX for comparison. In this assay, F9-K45Q-K77Q-ROX exhibited a longer half-life compared to 5-ROX alone, with t_1/2_ values of 17.9 and 4.7 min, respectively (Fig. 4A). In addition, we used a cohort of CD1 mice (n = 3, dose = 10 mg/kg) to evaluate the biodistribution of F9-K45Q-K77Q-ROX nanobody. At two hours post intravenous administration of the nanobody, we sacrificed the dosed animals and collected the brain, heart, lungs, spleen, kidneys, liver, leg muscle, and urine. Then, we imaged the fresh organ samples under an In Vivo Imaging System (IVIS) and measured the fluorescence intensity in each organ using ImageJ software. Evaluation of the biodistribution of F9-K45Q-K77Q-ROX demonstrated that the nanobody was primarily cleared by the kidneys (Figs. 4B and S5).

**Figure 4.**
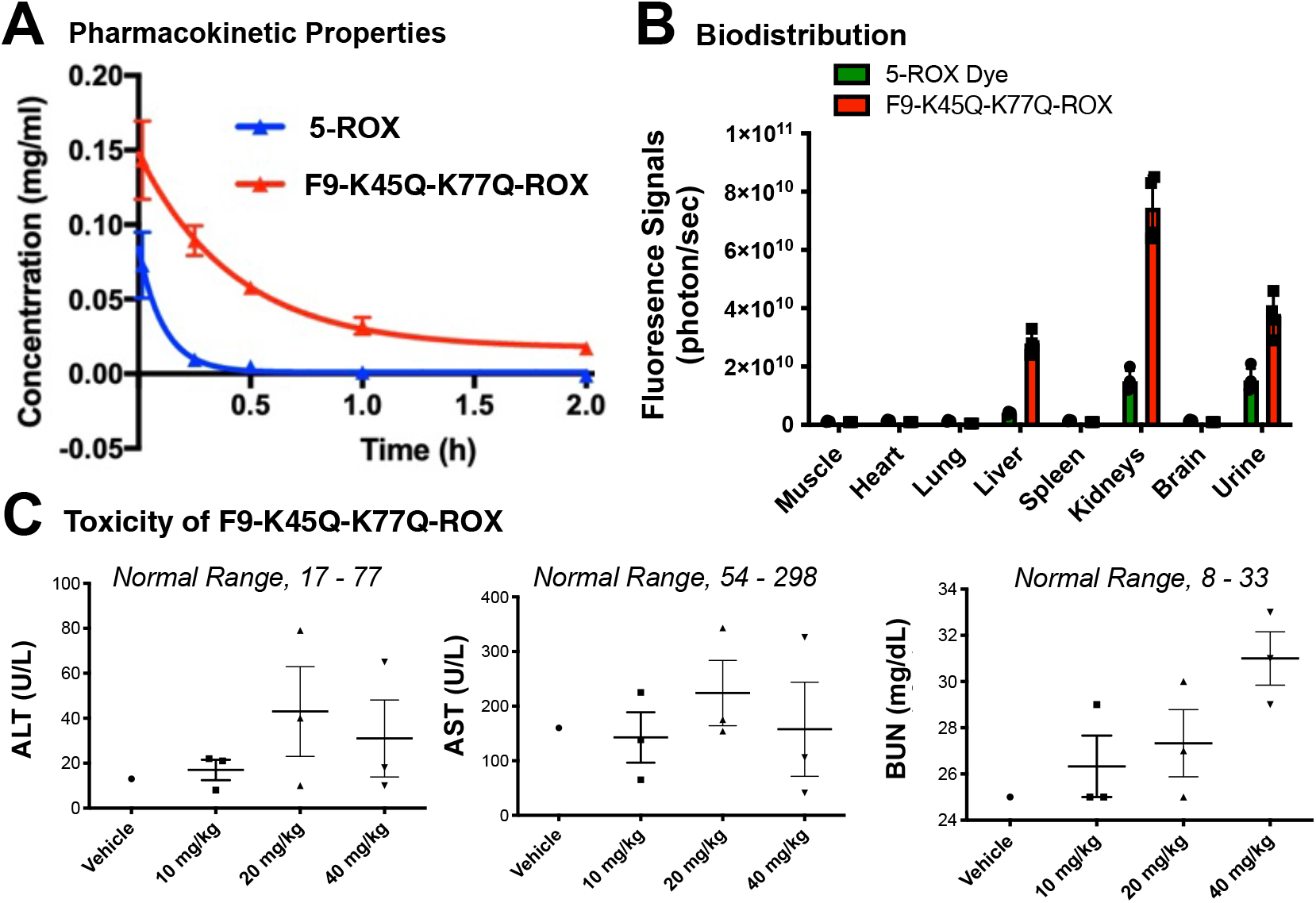
(A) In vivo pharmacokinetic properties of 5-ROX and F9-K45Q-K77Q-ROX in CD1 mice; (B) in vivo biodistribution of F9-K45Q-K77Q-ROX in CD1 mice; (C) serum markers of liver alanine aminotransferase (ALT), aspartate aminotransferase (AST), or blood urea nitrogen (BUN) at 24 h post intravenous administration of vehicle, 10 mg/kg, 20 mg/kg, or 40 mg/kg of F9-K45Q-K77Q-ROX.

*In vivo* toxicity of intravenously administered F9-K45Q-K77Q-ROX was tested in CD-1 mice using saline (vehicle) alone as the control. F9-K45Q-K77Q-ROX was injected at doses of 4 mg/kg, 20 mg/kg, or 40 mg/kg. Serum alanine aminotransferase (ALT), aspartate aminotransferase (AST), and blood urea nitrogen (BUN) were measured at 24 h post-injection. No increase in serum markers of liver or kidney toxicity was observed with F9-K45Q-K77Q-ROX-treated compared to saline-treated mice at any dose administered (Fig. 4C). Moreover, major organs were fixed immediately in 10% neutral-buffered formalin, embedded, sliced, and stained with hematoxylin and eosin (H&E) (Fig. S4). Slides were blindly reviewed by a board-certified veterinary pathologist for organ toxicity. No sign of toxicity was observed with F9-K45Q-K77Q-ROX within any treatment groups. The data further confirmed that the F9-K45Q-K77Q-ROX is non-toxic *in vivo* after a short-term exposure.

We developed azoxymethane (AOM)/dextran sodium sulfate (DSS)-induced colorectal adenomas in B6;129 mice (53). In this model, colorectal adenomas in the treated animals are typically observed at 8 weeks post-AOM administration. The AOM/DSS model recapitulates a multistep progression from colonic dysplasia to micro- and macro-adenomas, and ultimately carcinomas that mimics human colorectal cancer, although the genetic aberrations associated with the neoplasia are much more diverse than those found in human patients. AOM/DSS-induced tumors are located predominantly in the distal colon, which can be easily accessed by a colonoscope, and are macroscopically flat, nodular, or polypoid, coinciding with the morphological diversity of human colorectal tumors (54, 55) (Fig. 5A-E). We used an immunoblotting assay (Fig. 5F-G) and an immunofluorescence assay (Fig. 5H) with conventional monoclonal antibodies to evaluate COX-2 expression in the adenoma tissues. The results showed high COX-2 expression in AOM/DSS-induced mouse colorectal adenomas as compared to the adjacent normal colon.

**Figure 5.**
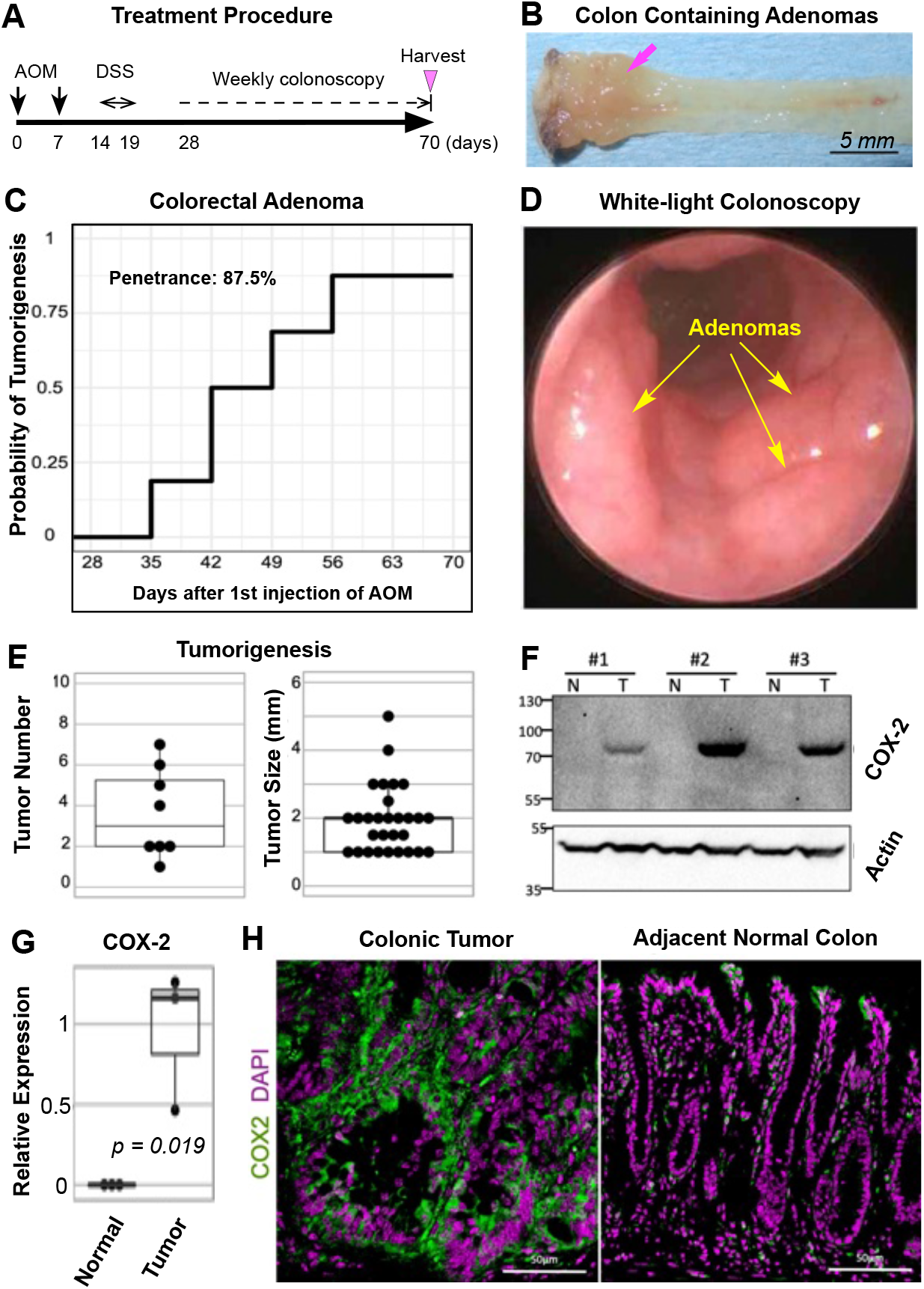
(A) Schematic of the AOM and DSS treatment procedure for the development of colorectal adenomas in B6;129 mice; (B) photograph of an excised colon containing adenomas; (C) tumor penetrance induced by AOM/DSS treatment; (D) white-light colonoscopy of adenomas at 10 weeks post-AOM/DSS treatment in B6;129 mice; (E) colon tumorigenesis in B6;129 mice at 10 weeks post-AOM/DSS treatment; (F) immunoblot assay showing COX-2 expression in colon adenomas versus normal colon tissues from AOM/DSS-treated B6;129 mice; (G) relative COX-2 expression in colon adenomas versus normal colon tissues; (H) immunofluorescence assay showing COX-2 expression in colon adenomas versus normal colon tissues from AOM/DSS-treated B6;129 mice.

We evaluated the ability of F9-K45Q-K77Q-ROX to target COX-2 and illuminate AOM/DSS-induced colorectal adenomas in B6;129 mice. In this experiment, B6;129 mice bearing colorectal adenomas were injected intravenously with F9-K45Q-K77Q-ROX (4 mg/kg) prior to fluorescence colonoscopy, enabling a clear delineation of colorectal adenomas at 30 min to 2 h post-administration (Fig. S5A). The imaging data showed a homogeneous distribution of the probe throughout the tumors with significantly higher tumor uptake than that of the surrounding normal tissues (Fig. 6A). The signal-to-noise ratios were determined by image analysis using ImageJ software as >12 (p < 0.001, n= 8 tumors) (Fig. 6B). For target validation, a blocking experiment was performed by intravenous dosing of a 10-fold excess of unlabeled F9-K45Q-K77Q nanobody at a 40 mg/kg dose 1 h prior to intravenous dosing of F9-K45Q-K77Q-ROX at 4 mg/kg dose, which resulted in minimal adenoma fluorescence (Figs. 6A,B & S6A,B). These data suggest that fluorescently labeled COX-2-targeted nanobodies can be useful as imaging agents for early colorectal cancer and potentially other neoplastic settings via endoscopy.

**Figure 6.**
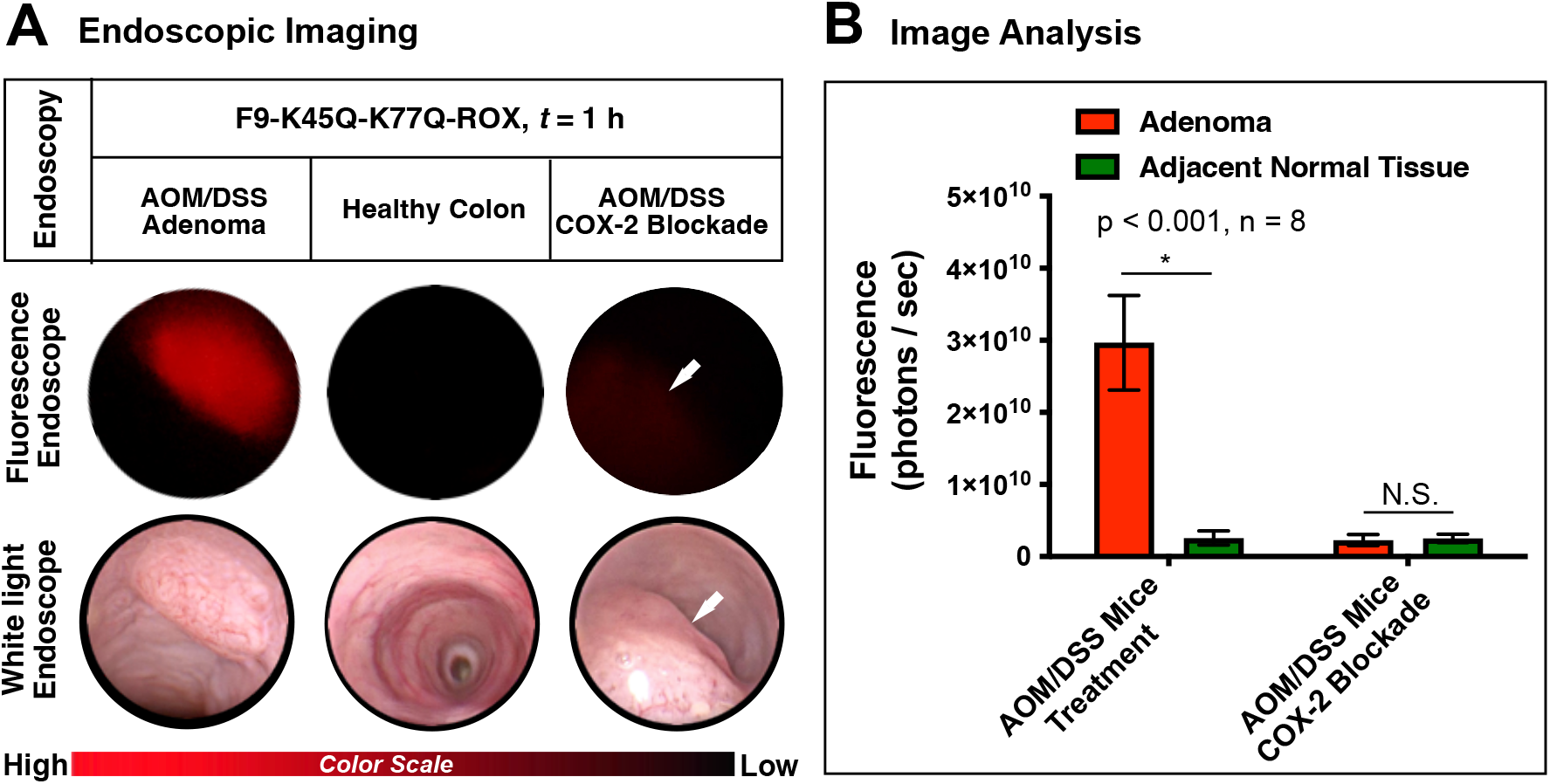
(A) Representative white-light and fluorescence in vivo colonoscopy images of B6;129 mice bearing AOM/DSS-induced colorectal adenomas or healthy colon dosed with F9-K45Q-K77Q-ROX nanobody (4 mg/kg, i.v.) or AOM/DSS-induced colorectal adenoma pre-treated with non-fluorescent F9-K45Q-K77Q (40 mg/kg, i.v.) 1 h prior to intravenous dosing of F9-K45Q-K77Q-ROX (4 mg/kg); (B) colonoscopy image analysis of colorectal adenomas of AOM/DSS B6;129 mice dosed with F9-K45Q-K77Q-ROX at 1 h post-injection or of AOM/DSS-induced colorectal adenoma pre-treated with non-fluorescent F9-K45Q-K77Q 1 h prior to dosing of F9-K45Q-K77Q-ROX using ImageJ software.

## Discussion

In this study, we identified and characterized F9, a novel single-domain antibody (nanobody) with high affinity and selectivity toward murine cyclooxygenase-2 (mCOX-2). The F9 nanobody was isolated through a phage display library constructed from the heavy-chain antibody repertoire of an alpaca immunized with native COX-2 protein, enabling the enrichment of fully functional VHH domains recognizing conformational epitopes on the enzyme. To elucidate the molecular determinants underlying F9–COX-2 recognition, we determined the crystal structure of the F9:COX-2 complex at high resolution. Structural analysis revealed that F9 engages one binding site per COX-2 monomer at the peroxidase active domain, forming an extensive interface dominated by hydrophobic and aromatic interactions with key residues that stabilize the complex. These contacts confer a sub-nanomolar to low-nanomolar dissociation constant (K_D), consistent with the strong binding affinity observed for the native nanobody in biophysical binding assays.

To facilitate site-specific labeling while preserving structural integrity, we introduced a double-point mutation (K45Q, K77Q) to eliminate two solvent-exposed lysine residues that could interfere with targeted fluorophore conjugation. The resulting F9-K45Q-K77Q mutant was subsequently modified with a carboxy-X-rhodamine (ROX) fluorophore through N-terminal amine conjugation. Proteomic characterization confirmed exclusive labeling at the N-terminus, validating the desired site-selectivity of the conjugation approach. Both the unlabeled F9-K45Q-K77Q and the fluorescent F9-K45Q-K77Q-ROX nanobodies demonstrated retained COX-2 binding affinity and selectivity relative to COX-1, mirroring the biochemical and structural properties of the parental F9 nanobody. Collectively, these results establish F9 as a robust molecular probe for COX-2, suitable for further engineering and imaging applications targeting inflammatory and neoplastic processes associated with COX-2 upregulation.

Fluorescence-based evaluation of the ROX-labeled analog F9-K45Q-K77Q-ROX was performed to assess its cell penetrance, intracellular localization, and binding selectivity toward cyclooxygenase-2 (COX-2) in head and neck squamous cell carcinoma (HNSCC) 1483 cells. Upon incubation with these cultured cells, confocal fluorescence microscopy revealed a distinct and reproducible pattern of red perinuclear fluorescence within the cytoplasmic compartment, suggesting that the probe not only effectively crossed the cellular membrane but also localized preferentially near perinuclear regions rich in endoplasmic reticulum, where COX-2 biosynthesis and retention predominantly occur. This intracellular distribution is consistent with efficient cellular uptake and high-affinity binding of F9-K45Q-K77Q-ROX to endogenous COX-2, supporting its suitability as a selective imaging probe for COX-2– expressing tumor cells. Beyond cell-based assays, pharmacokinetic profiling of F9-K45Q-K77Q-ROX in animal models demonstrated markedly improved in vivo stability and circulation time compared to the unconjugated ROX dye. The hybrid construct displayed an extended plasma half-life, reduced renal clearance, and enhanced tumor accumulation via the enhanced permeability and retention (EPR) effect characteristic of solid tumors. These favorable pharmacokinetic parameters—together with its robust fluorescent signal and selective intracellular binding—underscore the potential of F9-K45Q-K77Q-ROX as a dual-function imaging and diagnostic agent capable of delineating COX-2–expressing cancer lesions with high contrast and specificity in both in vitro and in vivo settings.

The F9-K45Q-K77Q-ROX conjugate was evaluated for its ability to efficiently penetrate cells and selectively recognize intracellular cyclooxygenase-2 (COX-2) using both fluorescence-based assays and live-cell imaging. In cultured 1483 human head and neck squamous cell carcinoma (HNSCC) cells, confocal fluorescence microscopy revealed distinct red perinuclear puncta corresponding to ROX emission, signifying effective cell uptake and intracellular localization of the probe. This red fluorescence pattern, concentrated in the perinuclear region, is characteristic of cytoplasmic distribution near the endoplasmic reticulum— consistent with the known subcellular localization of COX-2—thereby confirming the specific intracellular engagement of F9-K45Q-K77Q-ROX with its intended target. These findings demonstrate that the engineered mutations (K45Q and K77Q) did not negatively impact cellular permeability or target affinity but rather maintained or potentially enhanced the binding specificity toward COX-2 within tumor cells. In addition to its favorable cell-penetrating and target-binding properties, the ROX-labeled protein exhibited markedly improved in vivo pharmacokinetic behavior compared to the free dye. When administered systemically, F9-K45Q-K77Q-ROX displayed prolonged circulation and enhanced accumulation at tumor sites, consistent with passive tumor targeting through the enhanced permeability and retention (EPR) effect (56). Quantitative analysis revealed a plasma half-life on the order of *t*_*1/2*_ = 17.9 minutes, which represents an optimal balance for optical imaging applications—long enough to permit robust imaging of tumor tissue shortly after administration, yet sufficiently short to allow rapid systemic clearance and minimize background fluorescence in non-target organs. Collectively, these attributes underscore the potential utility of F9-K45Q-K77Q-ROX as an efficient and selective COX-2 imaging agent with desirable pharmacokinetic and optical characteristics for preclinical or translational imaging studies.

*In vivo* evaluation of the fluorescent nanobody conjugate F9-K45Q-K77Q-ROX revealed excellent biocompatibility and safety following a single systemic administration. Mice receiving the formulation exhibited no detectable signs of acute or delayed toxicity, as confirmed by routine clinical observation, histopathological inspection of major organs, and serum biochemistry analysis. These findings suggest that the amino acid substitutions (K45Q and K77Q) and ROX conjugation do not compromise the nanobody’s structural stability or induce undesirable immunogenic or cytotoxic effects in the biological system.

Functional imaging experiments performed by endoscopic visualization of colorectal adenomas demonstrated that the fluorescently labeled nanobody enabled vivid and sharply contrasted delineation of neoplastic lesions within the colon. Fluorescence signal accumulation at the adenoma sites produced distinct boundaries between tumor and normal mucosa, facilitating clear identification of the diseased tissue under minimally invasive imaging conditions. Specificity of the fluorescence labeling was rigorously validated by competitive blocking experiments in which pre-administration of excess unlabeled nanobody effectively abolished the fluorescent signal in the same regions, confirming that lesion-associated fluorescence arose from target-specific binding rather than nonspecific uptake or background accumulation. Together, these results establish F9-K45Q-K77Q-ROX as a highly selective, non-toxic optical probe suitable for real-time, targeted endoscopic imaging of colorectal adenomas in vivo. This nanobody-based system thus represents a powerful molecular tool for early lesion detection and potentially for guiding therapeutic intervention in gastrointestinal cancers.

Our laboratory and others have reported imaging probes for selective visualization of COX-2 in inflammation and cancer that provide a proof-of-principle for targeted detection of a wide range of neoplastic tissues that overexpress COX-2, including early CRC (32, 40, 57-62). The present fluorescent nanobody platform represents a transformative strategy for developing COX-2-selective imaging probes with substantial biochemical and translational advantages. First, the nanobody scaffold offers a remarkably facile mode of recombinant expression in various host systems, including bacterial, yeast, and mammalian cells, enabling efficient production at scale while maintaining structural fidelity. Second, the nanobody binds with high affinity to its target, cyclooxygenase-2 (COX-2), and exhibits pronounced selectivity over cyclooxygenase-1 (COX-1), a distinction crucial for minimizing off-target interactions and ensuring precise molecular imaging of inflammatory and neoplastic processes.

Third, its compact single-domain architecture tolerates extensive chemical or genetic manipulation—including site-specific conjugation of fluorophores—without significant loss of target-binding capability, thereby making it a versatile platform for multimodal probe engineering. In addition, this nanobody design displays notable physicochemical stability within the systemic circulation, resisting degradation and aggregation while maintaining structural integrity throughout the imaging window. This property enables broad biodistribution across tissues following intravenous administration, ensuring sufficient delivery to target sites. Importantly, the nanobody’s small size and inherent physicochemical properties confer superior cellular penetration capabilities, allowing the molecule to traverse cellular membranes efficiently and engage COX-2 localized within intracellular compartments—an advantage rarely achieved by larger antibody-based probes. Moreover, in vivo studies have demonstrated that the fluorescent COX-2 nanobody selectively accumulates in colorectal adenomas, enabling high-fidelity visualization of COX-2 expression patterns in diseased tissue. Following systemic administration, the probe yields images characterized by high tumor-to-background contrast, facilitating the noninvasive identification and delineation of early neoplastic lesions.

In conclusion, the results of these investigations establish a comprehensive foundation for the targeted use of COX-2–specific nanobodies as innovative molecular imaging agents for early colorectal cancer (CRC) detection through fluorescence endoscopy. By integrating molecular specificity with advanced optical imaging, this approach enables real-time, high-resolution visualization of COX-2 expression patterns at the mucosal surface, thereby facilitating identification of precancerous and early-stage neoplastic lesions that may otherwise evade conventional white-light endoscopy. The use of nanobody-based probes provides several unique advantages over traditional antibody or small-molecule techniques, including superior tissue penetration, rapid systemic clearance, high target-to-background contrast, and facile chemical conjugation to fluorescent dyes or radionuclides for multimodal imaging applications. Collectively, these attributes highlight the translational potential of this technology for precision-guided endoscopic diagnostics. While this study employed an early-stage murine model of CRC to validate molecular targeting and in vivo imaging performance, the versatility of the nanobody platform extends far beyond colorectal applications. Because COX-2 overexpression is a hallmark of numerous inflammation-driven cancers and precancerous conditions, this imaging strategy could be readily adapted to detect and monitor neoplastic diseases in other organ systems, including the esophagus, oral cavity, head and neck region, skin, and retina. In each of these tissues, COX-2 serves not only as an important biomarker of early tumorigenesis and disease progression but also as a potential indicator of therapeutic response to anti-inflammatory, chemopreventive, or targeted agents. Thus, implementing COX-2–targeted molecular imaging with nanobody probes could enable more personalized disease stratification, improved prognostication, and real-time assessment of treatment efficacy in both preclinical and clinical contexts.

## Materials and Methods

### Development of Anti-COX-2 Nanobodies

We developed COX-2-targeted nanobodies in collaboration with Turkey Creek Biotechnology, Waverly, TN, using established protocols (47, 48). An alpaca was immunized every 2 weeks for 12 weeks with 125 μg of purified COX-2 diluted 1:1 in Gerbu adjuvant (63). All animal procedures were performed at Turkey Creek Biotechnology (Waverly, TN) and strictly adhered to the United States Department of Agriculture Animal Welfare Act regulations for animal care and welfare. One-week following the last immunization, blood was drawn from the alpaca, and peripheral blood mononuclear cells (PBMCs) were isolated using SepMate™ tubes and density gradient centrifugation following the manufacturer’s protocol (STEMCELL Technologies). A cDNA library was made from the PBMC RNA by reverse transcription, and a nested PCR strategy was used to amplify coding regions of Vhh fragments as previously described (63). The resulting PCR fragments were cloned into a modified pADL22 vector (Antibody Design Labs) with hexahistidine and HA tags, and phage was produced using CM13 helper phage following the manufacturer’s protocols (Antibody Design Labs). A single round of panning was performed against murine COX-2 immobilized on a Maxisorb 96 well plate, and a promising clone, called “F9”, was isolated. In addition, the nanobody screening results showed no COX-1 binding.

### Nanobody Expression

The nanobody clone F9 was isolated from V_H_H sequences produced in a phage library against the antigen, mCOX-2. The gene of clone F9 in a pADL-22c phagemid vector was transformed into BL-21 (DE3) *E. coli* cells. A single bacterial colony containing the F9 gene was grown in 100 mL of starter culture (LB broth, 100 μg/mL ampicillin) at 37◦C, 225 rpm, overnight. Starter culture was then used to inoculate auto-induction growth medium containing glucose and α-lactose. The cells were grown at 37◦C, 225 rpm for approximately 5 h. The temperature was then lowered to 23◦C and shaking continued at 225 rpm for an additional 12 h. Following expression, cells were harvested by centrifugation.

### Site-directed Mutagenesis

The double mutant F9-K45Q-K77Q nanobody was generated using standard site-directed mutagenesis techniques (64). Briefly, the wild-type F9 nanobody in pADL-22c served as the template, and the following primers were used to make point mutations:

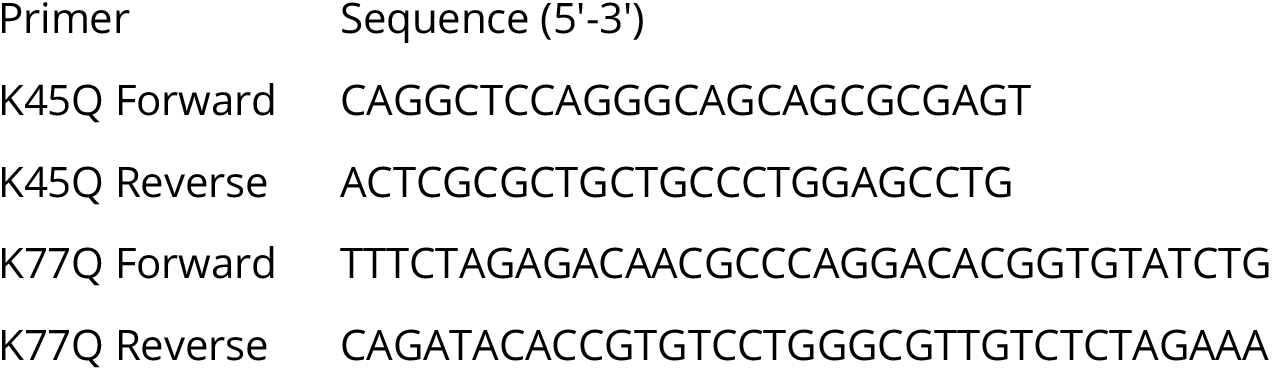

Standard PCR was performed using a Thermo Fisher ProFlex PCR System and PfuUltra High-Fidelity DNA polymerase (Agilent Technologies). Mutations were verified with Sanger sequencing (GenHunter, Nashville, TN).

### Nanobody Purification

The expressed nanobody was purified by resuspending the cell pellet in lysis buffer (10 mL/1 g cells) containing 5 M NaCl, 10% NP-40, 1 M Tris, and H_2_O at a ratio of 0.3:1.0:0.5:8.2 (v/v/v/v), and cells were homogenized by a PYREX® Dounce tissue grinder(65). Following centrifugation at 16,500 rpm for 30 min, cell lysate was incubated with Ni-NTA resin for at least 1 h. The lysate was added to a gravity column and purified with phosphate-buffered saline (PBS) (pH 7.4) containing 50 mM imidazole. Clone F9 was eluted from the Ni-NTA resin using buffer containing 250 mM imidazole. The Ni-NTA elution fractions were concentrated and then purified using size-exclusion chromatography (Superdex 75 16/600 column; GE Healthcare). Peak fractions were collected and tested for purity with SDS-PAGE. Pure fractions were collected and concentrated to approximately 10 mg/mL (1 mL) in PBS and stored at -80◦C. MALDI MS calcd for F9-K45Q-K77Q (M^+^) 15,143; found 15,143.

### Bio-conjugation Chemistry

The F9-K45Q-K77Q nanobody (2.5 mg in 250 mL, 0.00017 mmol) was diluted with sodium bicarbonate buffer (pH 9.2, 1.5 mL) and added dropwise to a freshly prepared solution of 5-carboxy-X-rhodamine N-succinimidyl ester (0.87 mg, 0.0014 mmol, 8 equiv.) in dimethyl sulfoxide (150 μL). The reaction mixture was gently stirred at 4 °C for 16 h. Following conjugation, excess unreacted dye was removed using a 10 kDa molecular weight cutoff centrifugal ultrafiltration spin column, followed by gel filtration. The reaction mixture was diluted sixfold with Tris buffer (20 mM Tris, 50 mM NaCl, 0.5 mM β-mercaptoethanol, 0.2 mM EDTA, 5% glycerol, pH 7.5) and centrifuged at 4000 rpm. The sample was concentrated to approximately 1 mL, and the filtrate was discarded. This dilution–centrifugation process was repeated with a 14-fold dilution in the same buffer until the filtrate became colorless. The concentrated sample (1 mL) was further purified by size-exclusion chromatography using a Superdex 200 prep grade column (GE Healthcare) on an FPLC system, with elution in Tris buffer. Fractions containing the conjugated nanobody were pooled and concentrated to 1 mL. Buffer exchange into PBS was performed using a 10 kDa MWCO centrifugal spin column, yielding a final concentration of 10 mg/mL (0.00017 mmol). The protein concentration was determined using the Pierce® BCA protein assay with bovine serum albumin as the standard. The purified F9-K45Q-K77Q-ROX conjugate was aliquoted in PBS and stored at −80 °C. The isolated yield of F9-K45Q-K77Q-ROX was 80%, corresponding to a final concentration of 2.1 mg/mL (1 mL) in PBS. MALDI MS calcd for F9-K45Q-K77Q-ROX (M^+^) 15,659; found 15,659.

### Degree of Labeling

The degree of labeling (DOL) was determined using a Tecan plate reader. 2 µl of F9-K45Q-K77Q or F9-K45Q-K77Q-ROX nanobody was deposited on each well, and absorption at 280 and 581 nm was measured. The DOL was calculated using the following two equations:

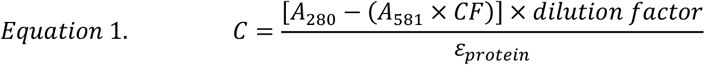

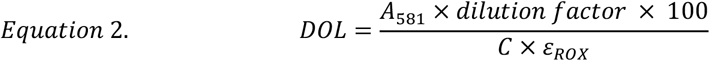

Where C is protein concentration (M), A_280_ and A_581_ are absorbance at 280 and 581 nm, respectively, CF is the correction factor for ROX dye (0.34), dilution factor is 10 or 100, ε_protein_ is the protein extinction coefficient (33,000 M^-1^cm^-1^), and ε_ROX_ is the ROX extinction coefficient (36,000 M^-1^cm^-1^). The DOL is expressed as mol dye/ mol protein. In a typical preparation, degree of labeling was calculated to be 99.55%.

### Proteomic Analysis of F9-K45Q-K77Q-ROX

Protein samples were combined with SDS-PAGE LDS sample buffer (3 mL 1 M Tris-HCl pH 7, 0.5 mL of double distilled water, 1.0 g SDS, 1 mL 0.1% bromophenol blue, 4 mL 100% glycerol, and 2 mL 15 M β-mercaptoethanol (100% stock) with 50 mM dithiothreitol) and were loaded onto a Novex 4-12% Bis-tris gel. The gel was stained with Novex colloidal Coomassie stain (Invitrogen), and de-stained in water. Gel bands were extracted, diced into 1 mm^3^ cubes, treated for 30 min with 45 mM dithiothreitol, and available Cys residues were carbamidomethylated with 100 mM iodoacetamide for 45 min. Gel pieces were destained with 50% acetonitrile in 25 mM ammonium bicarbonate, and proteins were digested with trypsin (10 ng/μL) in 25 mM ammonium bicarbonate overnight at 37 °C. Peptides were extracted by 60% acetonitrile, 0.1% trifluoracetic acid and 39.9% water. The extracts were dried by speed vac centrifugation and reconstituted in 0.1% formic acid. Peptides were then analyzed by LC-coupled tandem mass spectrometry (LC-MS/MS). First, an analytical column was packed with 20 cm of C18 reverse phase material (Jupiter, 3 μm beads, 300 Å, Phenomenex) directly into a laser-pulled emitter tip. Peptides were loaded on the capillary reverse phase analytical column (360 μm O.D. x 100 μm I.D.) using a Dionex Ultimate 3000 nanoLC and autosampler. The mobile phase solvents consisted of 0.1% formic acid, 99.9% water (solvent A) and 0.1% formic acid, 99.9% acetonitrile (solvent B). Peptides were gradient-eluted at a flow rate of 350 nL/min, using a 90-min gradient. The gradient consisted of the following: 1-72 min, 2-40% B; 72-78 min, 40-90% B; 78-79 min, 90% B; 79-80 min, 90-2% B; 80-90 min (column re-equilibration), 2% B. Upon gradient elution, peptides were analyzed using a data-dependent method on a Q Exactive Plus mass spectrometer (Thermo Scientific), equipped with a nanoelectrospray ionization source. The instrument method consisted of MS1 using an MS AGC target value of 3 × 10^6^, followed by up to 16 MS/MS scans of the most abundant ions detected in the preceding MS scan. The MS2 AGC target was set to 5 × 10^4^, dynamic exclusion was set to 10 s, HCD collision energy was set to 28 nce, and peptide match and isotope exclusion were enabled. For identification of peptides, tandem mass spectra were searched with Sequest (Thermo Fisher Scientific) against an *E. coli* database created from the UniprotKB protein database (www.uniprot.org), appended with the F9 nanobody sequence. Variable modification of +15.9949 on Met (oxidation), +57.0214 on Cys (carbamidomethylation), and +516.2049 (mass shift of 5-ROX) on Lys and the N-terminus of the protein were included for database searching. Search results were assembled using Scaffold 4.3.2. (Proteome Software).

### Fluorometry

Fluorescence excitation (λ_ex_) and emission (λ_em_) spectra of F9-K45Q-K77Q-ROX were recorded on a photon counting fluorometer operated with Felix software. The measurements were conducted in PBS at an optical density of 0.05/cm using a 10 × 4 mm cuvette at a 1 μM sample concentration. The fluorescence quantum yield (*φ*) of F9-K45Q-K77Q-ROX was calculated by comparing the fluorescence intensity with the fluorescence intensity of 5-ROX. The extinction coefficient (ε) of F9-K45Q-K77Q-ROX was calculated using the absorbance spectra of solutions (1 μM to 12 μM) measured in 10-mm pathlength cuvettes recorded with a UV–VIS spectrophotometer (UV-2501PC, Shimadzu Corporation, USA).

### *In Vitro* COX-2:Nanobody Binding

We used a Monolith NT.115 (green/blue) microscale thermophoresis (MST) instrument (NanoTemper Technologies) to determine the COX-2 binding K_D_ (dissociation constant) values of our nanobodies. We prepared serial dilutions of nanobodies dissolved in PBS buffer at pH 7.4. Purified mCOX-2 (15 μM, PBS pH 7.4) was fluorescently labelled with Atto665 N-succinimidyl ester (Sigma-Aldrich). The COX-2 enzyme in PBS was diluted in sodium bicarbonate buffer (pH 9.2), which was added to Atto665 N-succinimidyl ester solution in methyl sulfoxide and stirred overnight gently at 4°C. Excess free dye was removed by repetitive ultrafiltration in a centrifugal spin column (molecular weight cut off: 50 kDa). The Atto665-labeled COX-2 (5 nM) was then brought into PBS pH 7.4 and incubated for 5 min with serial dilutions of the test nanobodies in PBS, pH 7.4. Measurements were performed using 20% and 40% MST power and between 20–80% LED power at 24 °C. MST traces were recorded using 5 s MST power off, 30 s MST power on, and 5 s MST power off. The program software plots the thermophoresis signal against the concentration of the ligand to generate a binding curve, which allows calculation of the dissociation constant (K_D_) values.

### *In Vitro* COX Inhibition Assay

Purified ovine COX-1 and murine COX-2 isozyme inhibitory activities of the nanobodies were evaluated by modification of methods reported by Mitchener et. al. (66). Five solutions were prepared: 1) apoenzyme in Tris-HCl (pH 8.0 with 0.5 mM phenol); 2) substrate [AA plus 5-phenyl-4-pentenyl-1-hydroperoxide (PPHP) in DMSO]; 3) F9, F9-K45Q-K77Q or F9-K45Q-K77Q-ROX in PBS; 4) heme in Tris-HCl; and 5) quench (0.5 % acetic acid in ethyl acetate). The final concentrations in the incubation mixtures were 15 nM oCOX-1 or mCOX-2,10 µM AA, 50 nM PPHP, 30 nM heme, and variable concentrations (500, 250, 125…. 3.915, 0 nM) of test nanobodies. The final constitution of the reaction mixture medium was 93.3% Tris-HCl, 3.3% PBS, and 3.3% DMSO (pH 7.4) before the addition of quench solution. Under standard assay conditions, enzyme and heme were combined prior to the addition of F9, F9-K45Q-K77Q, or F9-K45Q-K77Q-ROX nanobody at the desired concentration. Following incubation at 25 °C for 10 min, and then 37 °C for 5 min, substrate solution containing AA and PPHP (which is added to eliminate a kinetic lag phase) was added. Alternatively, F9, F9-K45Q-K77Q, or F9-K45Q-K77Q-ROX nanobody at the desired concentration was added to apoenzyme solution and allowed to pre-incubate at 25 °C for 15 min. Next, heme solution was added followed by incubation for another 15 min at 25 °C and then 3 min at 37 °C. Then, substrate solution was added. In both cases, the final mixture was allowed to react at 37 °C for 10 s before combining with an equivalent volume of quench solution with internal standards for PGE_2_/D_2_ (15 nM). The short reaction time was employed to minimize substrate consumption and reaction-dependent enzyme inactivation. The organic phases were removed, dried under N_2_ gas, reconstituted in 1:1 methanol: H_2_O, and chromatographed using reverse-phase HPLC on a *Luna* C18 (2) column (50 × 2 mm, 5 μm) (Phenomenex, Torrance, CA) (solvent A: 5 mM ammonium acetate, pH 3.6; solvent B: acetonitrile with 6% solvent A) in line with a 3200Q trap tandem mass spectrometer (AB SCIEX) operated in positive ion mode. Samples were monitored for PGE_2_ (*m/z* 370.2→317.2), PGD_2_ (*m/z* 370.2→317.2), and the respective internal standard on an elution gradient of 20-90% solvent B over 5 min followed by 90% solvent B for 1.4 min at a flow rate of 300 μL/min. The data were processed with Analyst software.

### X-Ray Crystallography

We co-crystallized the F9 nanobody with purified apo mCOX-2 using a drop vapor diffusion method as described previously (52). A solution of purified apo mCOX-2 (11.5 mg/mL) was mixed with a 2-fold molar excess of F9 in PBS. The mixture was incubated at 25°C for 2-3 weeks for the crystals to form and grow to their full size. Then, the crystals were mounted and flash frozen in liquid nitrogen for shipment and data collection at the Northeastern Collaborative Access Team beam line 24-ID-C at the Advance Photon Source at the Argonne National Laboratory. We processed the diffraction data with XDS (67), revealing compound F9:COX-2 cocrystals belonging to space group *P*2_1_2_1_2. We used a search model (PDB id: 3NT1) and determined the structure by molecular replacement using Phaser (68) Structure refinement was done in PHENIX (PMID 12393927)(69), The resulting atomic coordinates and structure factors have been deposited to the Protein Data Bank (PDB) at http://www.rcsb.org. The PDB access id for the F9:COX-2 structure complex is 8ET0.

### *In Vitro* Cell Imaging

Fluorescence imaging of COX-2 in live cells by F9-K45Q-K77Q-ROX was performed using both 1483 HNSCC cells (COX-2-expressing) and OVCAR3 cells (COX-1-expressing). Cells were grown to 50-60% confluency in MatTek dishes on glass inserts in 2.0 mL DMEM/F12 medium + 0.1% fetal bovine serum (FBS) (1483 cells) or 2.0 mL RPMI 1640 + 0.1% FBS (OVCAR3 cells). F9-K45Q-K77Q-ROX [100 nM] was added to the cells for 60 min at 37°C. The cells were then washed 3 times with Hanks balanced salt solution (HBSS), followed by imaging in 2.0 mL fresh HBSS/Tyrode’s on a Leica DM IL LED Fluorescent Microscope using a Texas Red filter (49008-ET) with 50 ns exposure. Overlays were performed with fluorescence and bright field to demonstrate subcellular localization of signal. All treatments were performed in duplicate dishes in at least three separate experiments. To block the F9 binding site, the cells were preincubated with 5 to 10 μM unlabeled nanobody for 30 min prior to the addition of F9-K45Q-K77Q-ROX.

### *In Vivo* Pharmacokinetic Assay

Male, 12-week-old CD-1 mice were housed with free access to food and water on a 12-h light/dark cycle. F9-K45Q-K77Q-ROX (100 µl, 290 µg) or ROX dye (100 µl, 300 µg) was injected intravenously via the tail vein (3 mice/treatment). Approximately 10 µl of whole blood was collected from the tail vein at 1, 15, and 30 min and at 1, 2, 4, and 8 h after injection. Blood was immediately plated in quadruplicate on a black NanoQuant plate (2 µl/well), and fluorescence was measured at excitation/emission 581/605 nm. Standard curves of F9-K45Q-K77Q-ROX or 5-ROX dye were prepared in whole blood. Circulation half-life was calculated in GraphPad Prism using a one-phase decay model.

### *In Vivo* Biodistribution Assay

*In vivo* biodistribution of F9-K45Q-K77Q-ROX was measured in male CD1 mice. Two hours after tail vein injection (100 µl, 290 µg), the mice were sacrificed, and the following organs were collected: brain, heart, lungs, spleen, kidneys, liver, leg muscle, and urine. The organs were imaged using an *In Vivo* Imaging System (IVIS) for nanobody fluorescence. Radiant efficiency ([p/s]/[µW/cm^2^]) was quantified for each organ.

### *In Vivo* Toxicity Assay

Saline or F9-K45Q-K77Q-ROX (doses 4, 20, and 40 mg/kg) was administered to wild-type CD-1 mice (4-6 weeks old) via tail vein injection. Blood, liver, and kidneys were collected at 24 h post-injection. Whole blood was incubated at room temperature for 30 min and centrifuged at 4,000 rpm for 5 min to obtain serum. Serum alanine aminotransferase (ALT), aspartate aminotransferase (AST), and blood urea nitrogen (BUN) were determined at the Vanderbilt Translational Pathology Shared Resource (TPSR) core facility using a commercially available Transaminase-CII kit and Blood Urea Nitrogen Test Wako, respectively. Liver and kidneys were fixed immediately in 10% neutral-buffered formalin, embedded, sliced 5 μm thick, and stained with hematoxylin and eosin. A board-certified pathologist at the TPSR examined the slides for organ toxicity.

### Animal Model of Colorectal Adenoma

All mice were housed, and experiments were conducted in accordance with a protocol approved by the Institutional Animal Care and Use Committee at Vanderbilt University Medical Center. To develop the animal model of colorectal adenomas, a procedure was used that we published recently (62). Briefly, we treated B6;129 mice with 10 mg/kg of azoxymethane (AOM; Sigma-Aldrich, St. Louis, MO) in PBS followed by free access to drinking water containing colitis-grade dextran sulfate sodium (DSS; #DS1044, Gojira Fine Chemicals, OH) at a concentration of 0.1 mg/mL. The mice were euthanized by carbon dioxide and subsequent cervical dislocation at the time of tissue harvesting or when they met one of the following humane end points; 20% of body weight loss from baseline, severe rectal prolapse, or severe bleeding due to tumor development.

### Immunofluorescence Assay

Mouse tissues were freshly harvested after appropriate euthanasia, immediately fixed with 4% paraformaldehyde in PBS at 4 °C for 4 h, and submerged into 30% sucrose at 4 °C, overnight. Specimens were embedded into O.C.T. compound (Sakura Finetek, Torrance, CA), and then five-micrometer cryosections were prepared for immunofluorescence. The sections were blocked with Abcam protein block (Abcam, Ab156024, Cambridge, MA) for 5 min, and incubated with anti-COX2 antibody (Cayman chemical, #160126, 1:200) at 4 °C, overnight. Donkey anti-rabbit IgG, Alexa 568 was used as the secondary antibody, and DAPI was used for nuclear staining. All imaging was done with a Nikon A1R laser confocal microscope.

### Immunoblotting Assay

Freshly harvested mouse colon and colonic tumors in RIPA buffer (10mM Tris-HCl, pH 8.0,1mM ethylenediaminetetraacetic acid, 0.5mM ethylene glycol-bis(β-aminoethyl ether)-N,N,N′,N′-tetraacetic acid, 1% Triton X-100, 0.1% Sodium Deoxycholate, and 0.1% Sodium dodecyl sulfate) with protease and phosphatase inhibitors were lysed by sonication at 4 °C. Lysates were boiled in LDS Sample Buffer (Novex, Carlsbad, CA) and used for SDS-PAGE and immunoblotting. Blots were incubated in 5% milk with 0.1% Tween-20 in Tris-buffered saline (TBS) for an hour at 22 °C. Membranes were incubated with anti-COX2 antibody (Cayman chemical, #160126, 1:200) at 4 °C overnight followed by an horseradish peroxidase-conjugated secondary antibody for an hour at 22 °C. Chemiluminescence signals were read by a Bio-Rad ChemiDoc XRS Imaging System, and quantification of protein bands was conducted using a Bio-Rad Image Lab software.

### *In Vivo* Endoscopic Imaging

Colonoscopies were performed with a Karl Storz Endoscope system (Karl Storz, Tuttlingen, Germany) under appropriate anesthesia with isoflurane (Piramal Critical Care Inc., Bethlehem, PA). The instrument was configured with a 1.9-mm, 30-degree, 10-cm rigid telescope attached to a tricam camera and a modified D-light source. The endoscopic system included an RFP/mCherry filter (Karl Strorz, #60100036), enabling simultaneous capturing of fluorescence and visible light still and video images using an Aida DVD-M camera while projecting endoscopic images on a TV monitor.

### Statistical Methods

We used the student’s t-test to compare treatment groups for statistical analyses. Statistical significance was set at a value of *P* ≤ 0.05, where the data were used as the arithmetic mean and standard error within the size of the samples (n).

## Acknowledgments

We thank the Vanderbilt Translational Pathology Shared Resource (TPSR) for blood chemistry and histopathological analyses of tissue samples, the Vanderbilt Cell Imaging Shared Resource for fluorescence microscopy, and the Vanderbilt University Institute of Imaging Science for *ex vivo* organ imaging. We also acknowledge the Vanderbilt Mass Spectrometry Research Center for ESI-MS and MALDI-MS analyses, and Dr. Carol Rouzer of the Vanderbilt University School of Medicine Basic Sciences for her critical reading and editorial assistance. This work was supported by the Vanderbilt Institute for Clinical and Translational Research (Grant Nos. VR54033 and VR54033.1 to M.J.U.), the Phi Beta Psi Sorority Trust (Grant No. AWD00000652 to M.J.U.), the National Cancer Institute (Grant Nos. R01 CA89450 to L.J.M.; R01 CA260958 to M.J.U. and C.L.D.; and R35 CA197570, UG3 CA241685, P01 CA229123, and P50 CA236733 to R.J.C.), and Faculty Incentive Awards from the Vanderbilt University School of Medicine Basic Sciences (Fund Nos. FF_300455 to M.J.U. and FF_242218 to L.J.M.).

## Author Contributions

M.J.U. designed the nanobody mutants and performed protein chemistry for the synthesis of fluorescent nanobody conjugates, determined photophysical properties and COX-2:nanobody dissociation constants, carried out *in vivo* pharmacokinetic and toxicity assays, executed *in vivo* endoscopic imaging studies, analyzed and interpreted the acquired data, and drafted the manuscript; S.X. performed wild-type F9 nanobody expression, purification, and wild-type F9:COX-2 co-crystallization; M.C.G. and A.M.A. performed expression and purification of F9-K45Q-K77Q and F9-K45Q-K77Q-ROX; H.N. developed the colonic adenoma animal model and performed endoscopic imaging experiments with M.J.U. and characterized COX-2 expression in colonic adenomas using immunoblotting and immunofluorescence assays. K.L.R. performed proteomic analysis of F9-K45Q-K77Q-ROX; B.C.C. performed cell-based fluorescence microscopy; S.B. conducted X-ray crystal diffraction and data collection; C.R.D. and E.L.R performed pharmacokinetic and organ distribution studies with M.J.U.; P.J.K. and J.K performed HPLC analysis of the nanobodies. M.L.R performed the MALDI-MS analysis; M.M and S.L. performed *in vitro* purified oCOX-1 and mCOX-2 inhibition assays; B.E.W. directed alpaca immunization and isolation of nanobody clones; B.W.S. directed cDNA synthesis, and phage display for nanobody clones; C.L.D. provided animals and instruments for pharmacokinetic and biodistribution studies; R.J.C. provided support for the animal model of colorectal adenoma; L.J.M. oversaw the overall program, designed and interpreted experiments, edited the manuscript and provided financial support for the study.

## Supporting Information

### HPLC Conditions

Solvent system, A: 10 mM ammonium acetate, pH’d to 3.1 with TFA, B: ACN + 10 mM ammonium acetate; 0.075% TFA and 10% H_2_O, Column: pentafluoro-propyl, 25 × 0.46cm @ 42C, gradient elution, Detection UV @ 235 nm and 278 nm (local max of F9-K45Q-K77Q nanobody), Fluorescence: ex = 581 nm & em = 610 nm.

**Figure S1.**
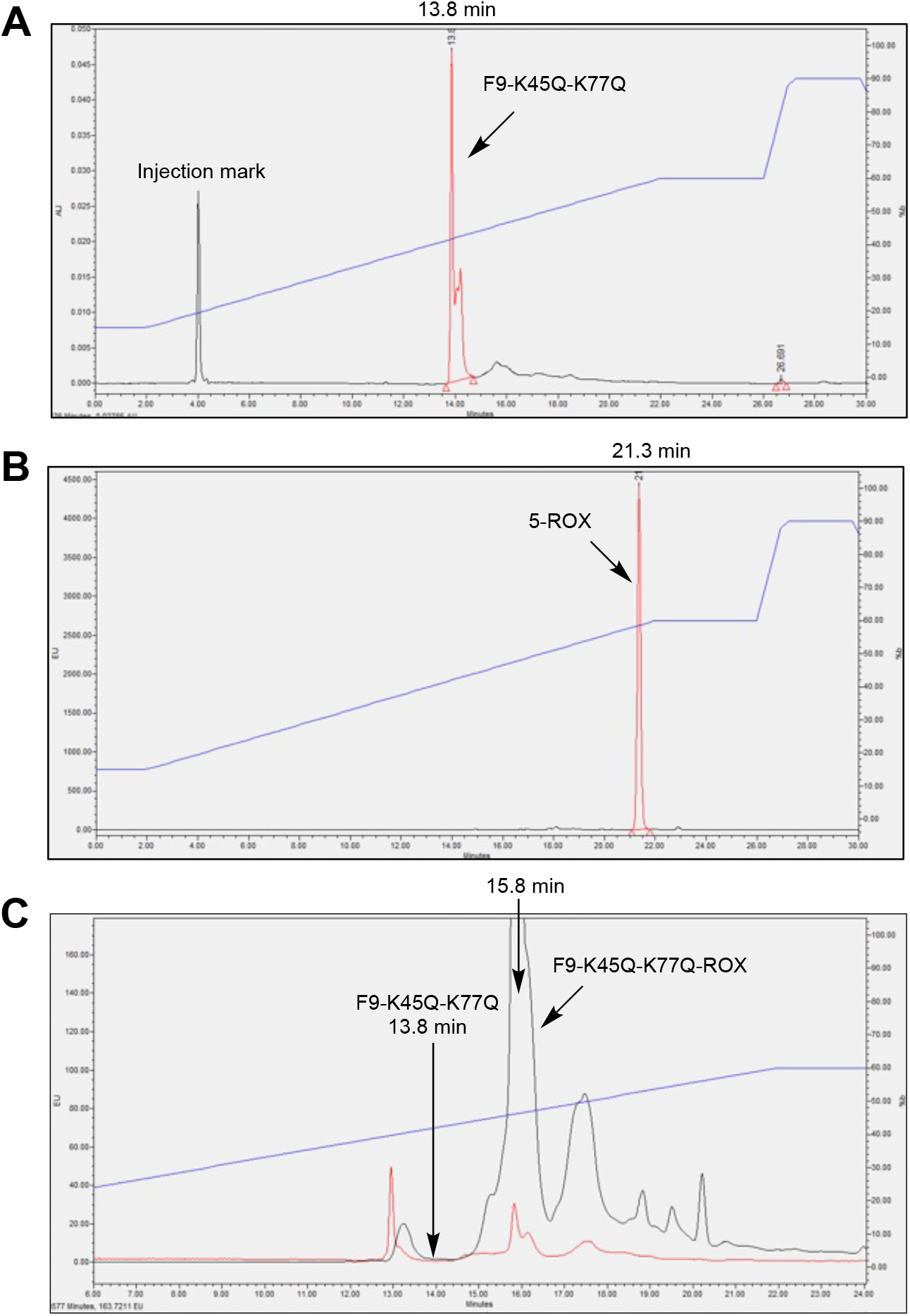
HPLC chromatography. (A) HPLC chromatogram of F9-K45Q-K77Q-ROX nanobody on VU (278 nm) detection, (B) HPLC chromatogram of 5-ROX dye on fluorescence (em 610 nm) detection, and (C) Overlay of UV (278 nm) and fluorescence (em 610 nm) HPLC chromatogram of the reaction mixture.

**Figure S2.**
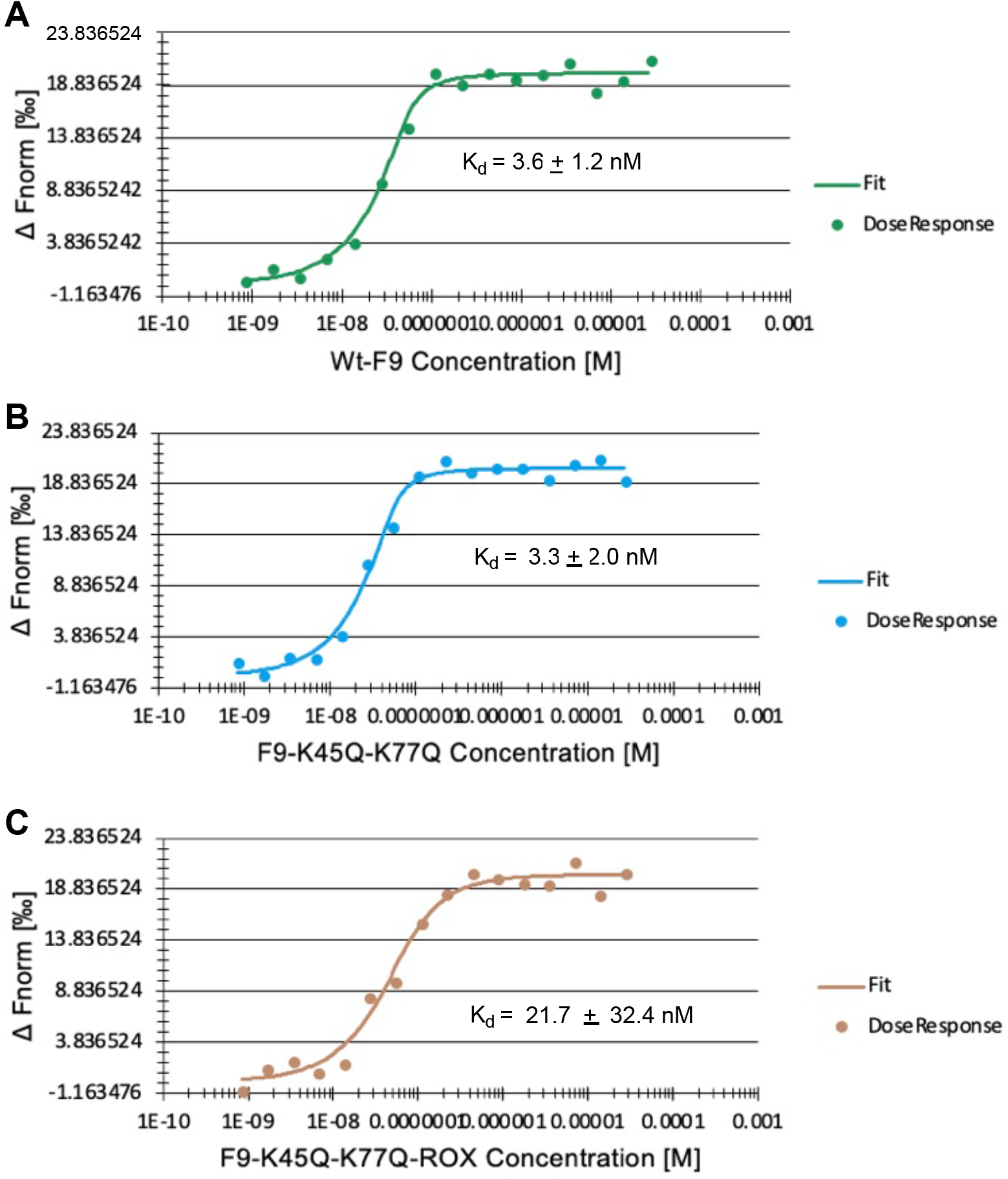
Microscale Thermophoresis (MST) assay determining COX-2:nanobody dissociation constant (K_*d*_) values on a NanoTemper Monolith NT.115 Instrument. (A) Purified COX-2:Wt-F9 binding curve, (B) purified COX-2:F9-K45Q-K77Q binding curve, (C) purified COX-2:F9-K45Q-K77Q-ROX binding curve.

**Figure S3.**
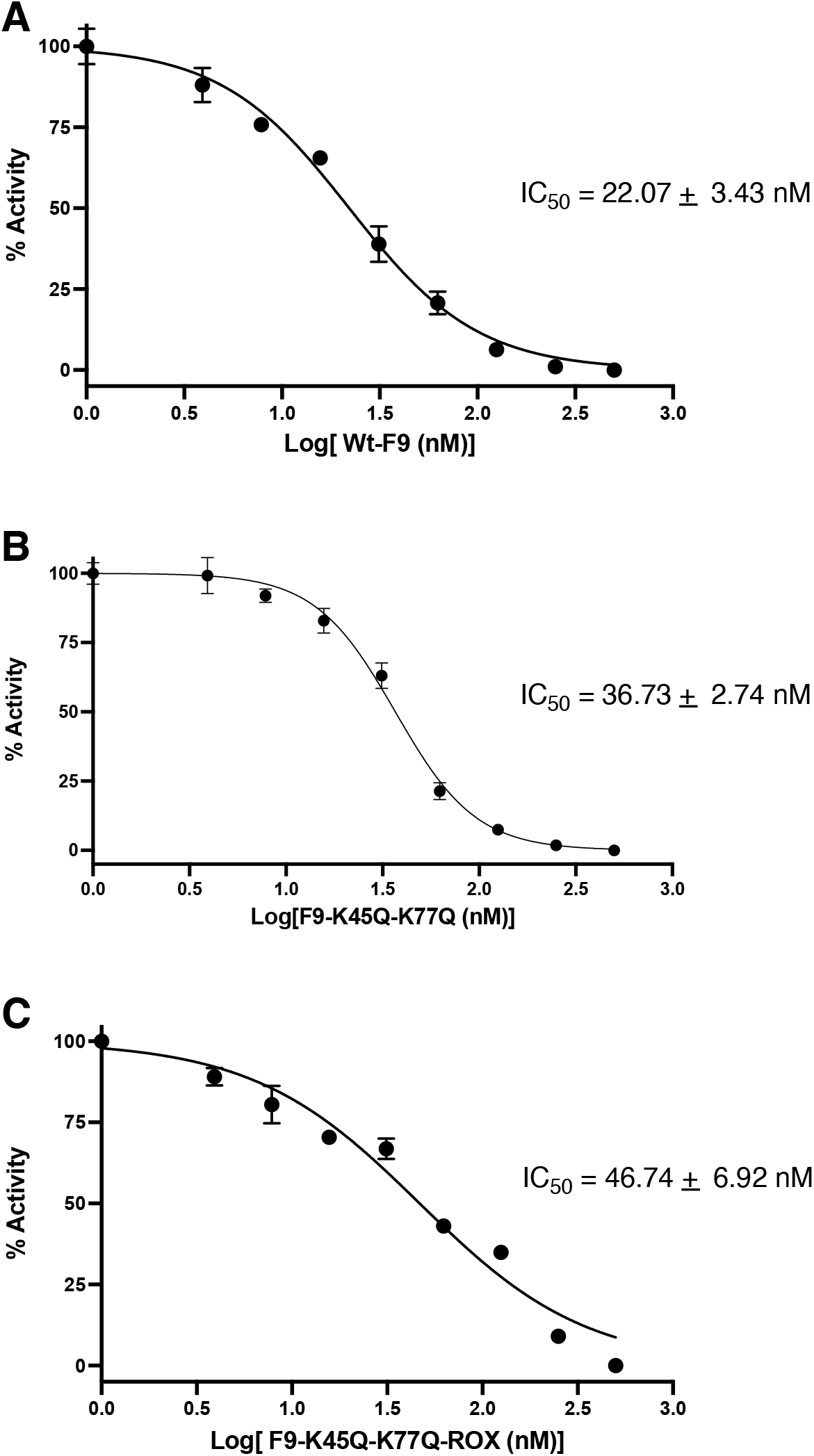
Inhibition of purified mCOX-2 by targeted nanobodies. (A) Dose-dependent inhibition of purified mCOX-2 by the Wt-F9 nanobody, (B) dose-dependent inhibition of purified mCOX-2 by the F9-K45Q-K77Q nanobody, (C) dose-dependent inhibition of purified mCOX-2 by the F9-K45Q-K77Q-ROX nanobody,

**Figure S4.**
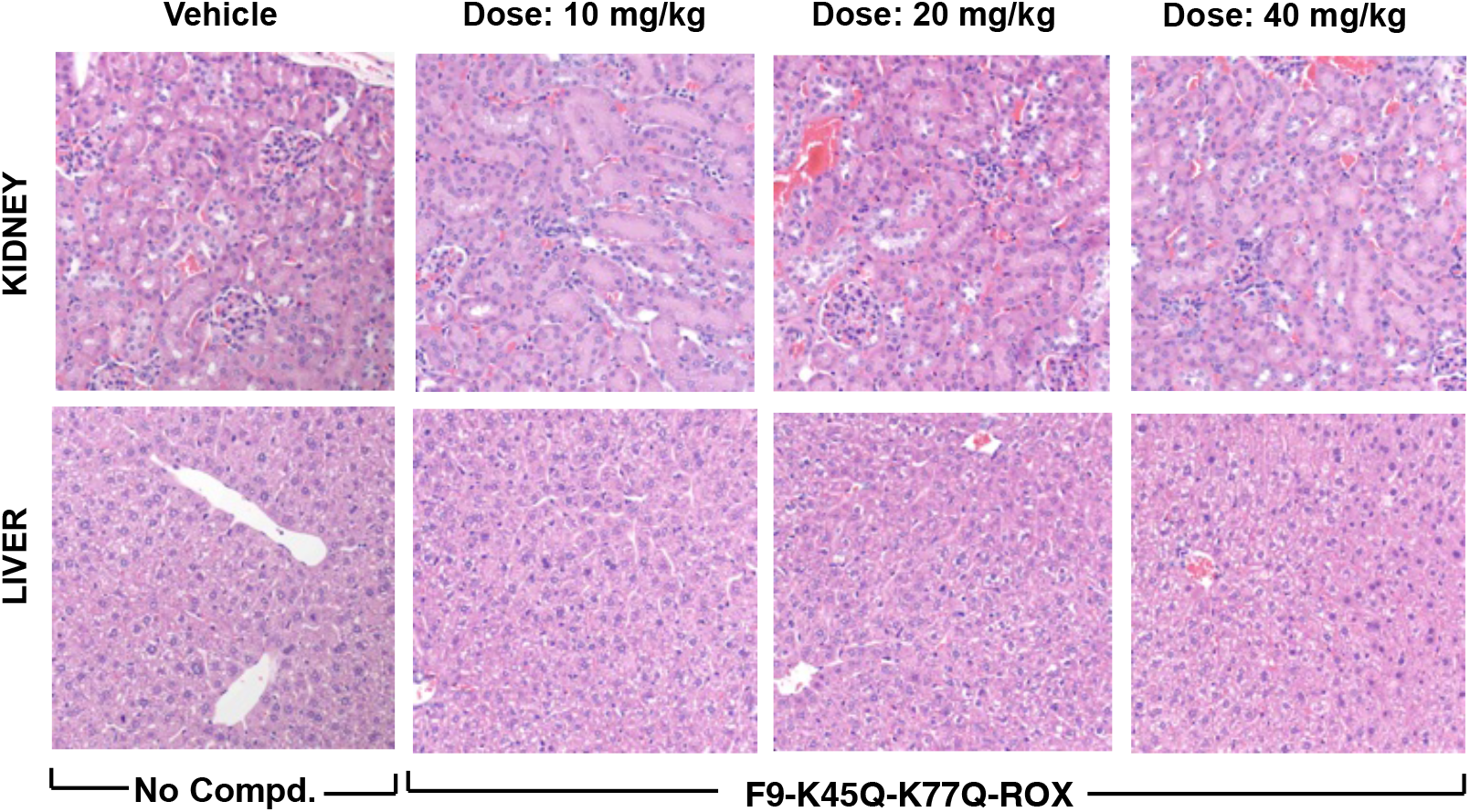
*In vivo* toxicity assay. H&E sections of liver and kidney at 24 h post intravenous administration of vehicle (PBS pH 7.4), 10 mg/kg, 20 mg/kg of F9-K45Q-K77Q-ROX, or 40 mg/kg of F9-K45Q-K77Q-ROX.

**Figure S5.**
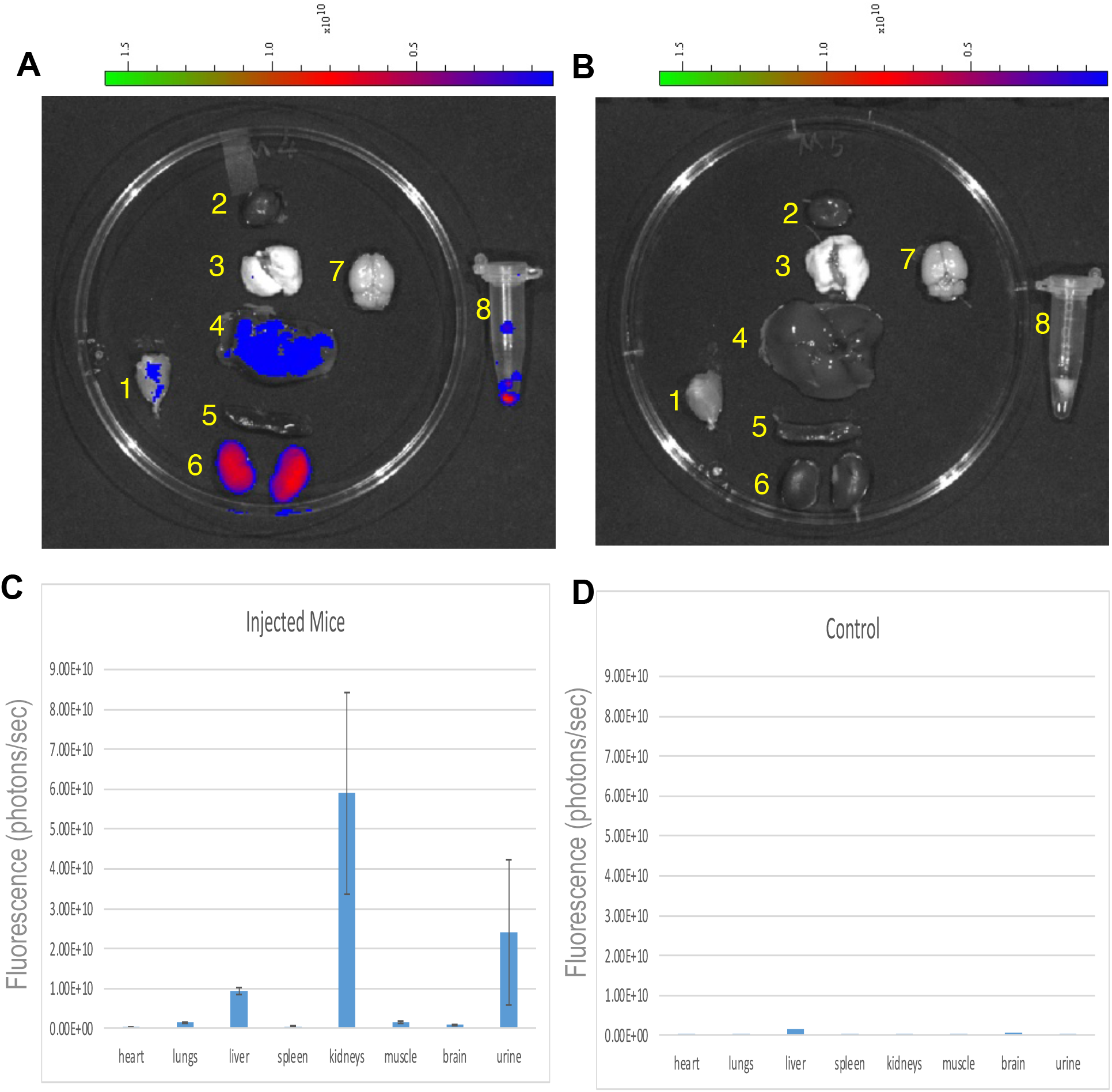
*Ex vivo* Imaging of organ distributions. (A) *Ex vivo* fluorescence image of the major organs (1. muscle, 2. heart, 3. lung, 4. liver, 5. spleen, 6. kidney, 7. brain, 8. Urine) of a CD1 mouse treated with F9-K45Q-K77Q-ROX (4 mg/kg, i.v., t = 2 h), (B) *ex vivo* fluorescence image of the major organs (1. muscle, 2. heart, 3. lung, 4. liver, 5. spleen, 6. kidney, 7. brain, 8. urine) of a control CD1 mouse treated with vehicle (PBS), (C) measurements of fluorescence signal in major organs (1. muscle, 2. heart, 3. lung, 4. liver, 5. spleen, 6. kidney, 7. brain, 8. urine) from an F9-K45Q-K77Q-ROX-injected mouse, or (D) vehicle-injected mouse using ImageJ software.

**Figure S6.**
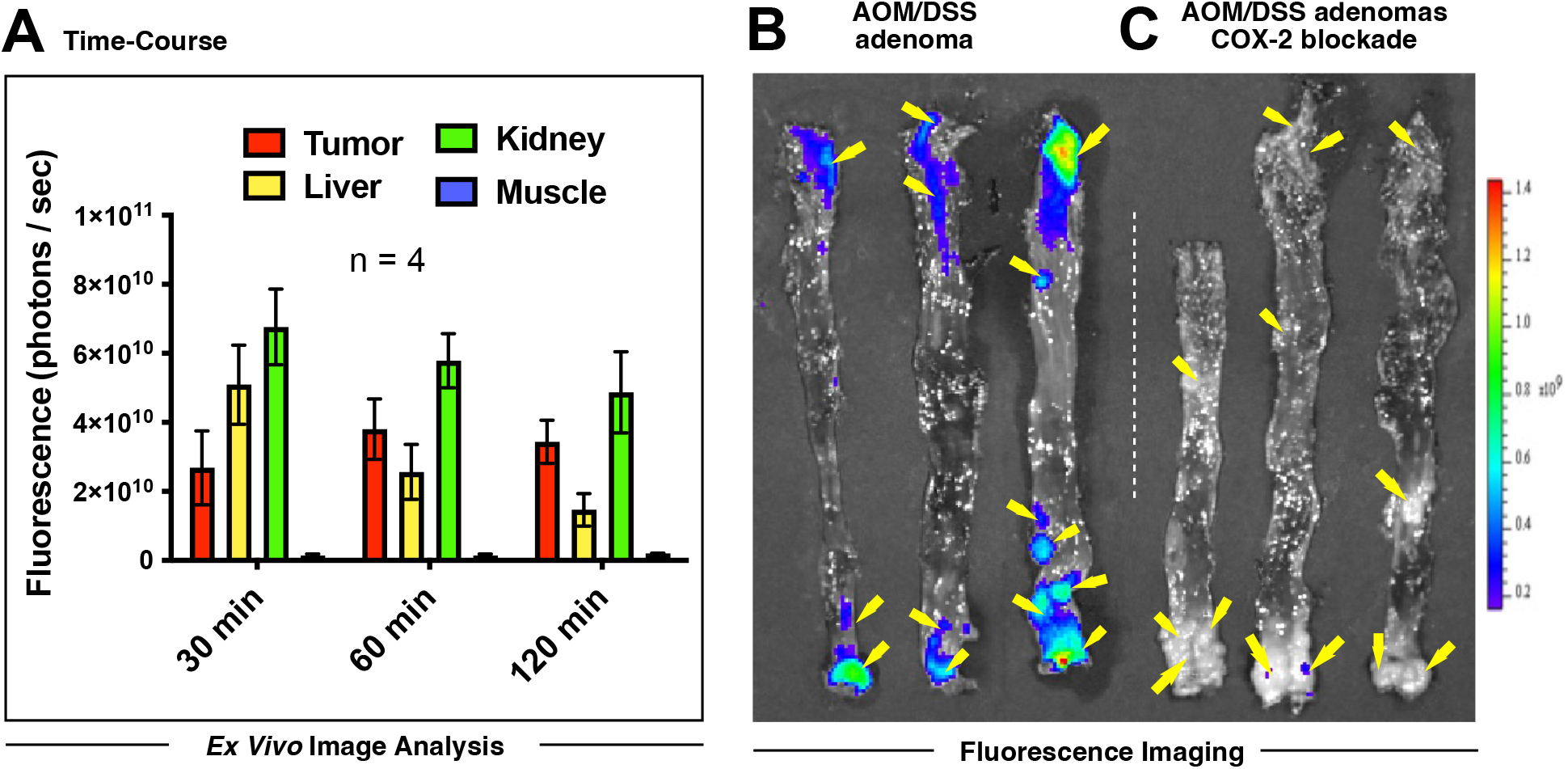
(A) Analysis of *ex vivo* organ images of B6;129 mice bearing AOM/DSS adenomas treated with F9-K45Q-K77Q-ROX (4 mg/kg, i.v.) at *t* = 0.5 h, 1 h, and 2 h post-injection; (B) *ex vivo* fluorescence image of colon tissues of B6;129 mice t bearing AOM/DSS adenomas treated with F9-K45Q-K77Q-ROX (4 mg/kg, i.v. t = 1 h); (C) *ex vivo* fluorescence image of colon tissues of B6;129 mice bearing AOM/DSS adenomas treated with F9-K45Q-K77Q (40 mg/kg, i.v.) 1 h prior to the injection of F9-K45Q-K77Q-ROX (4 mg/kg, i.v. t = 1 h).

**Figure S7.**
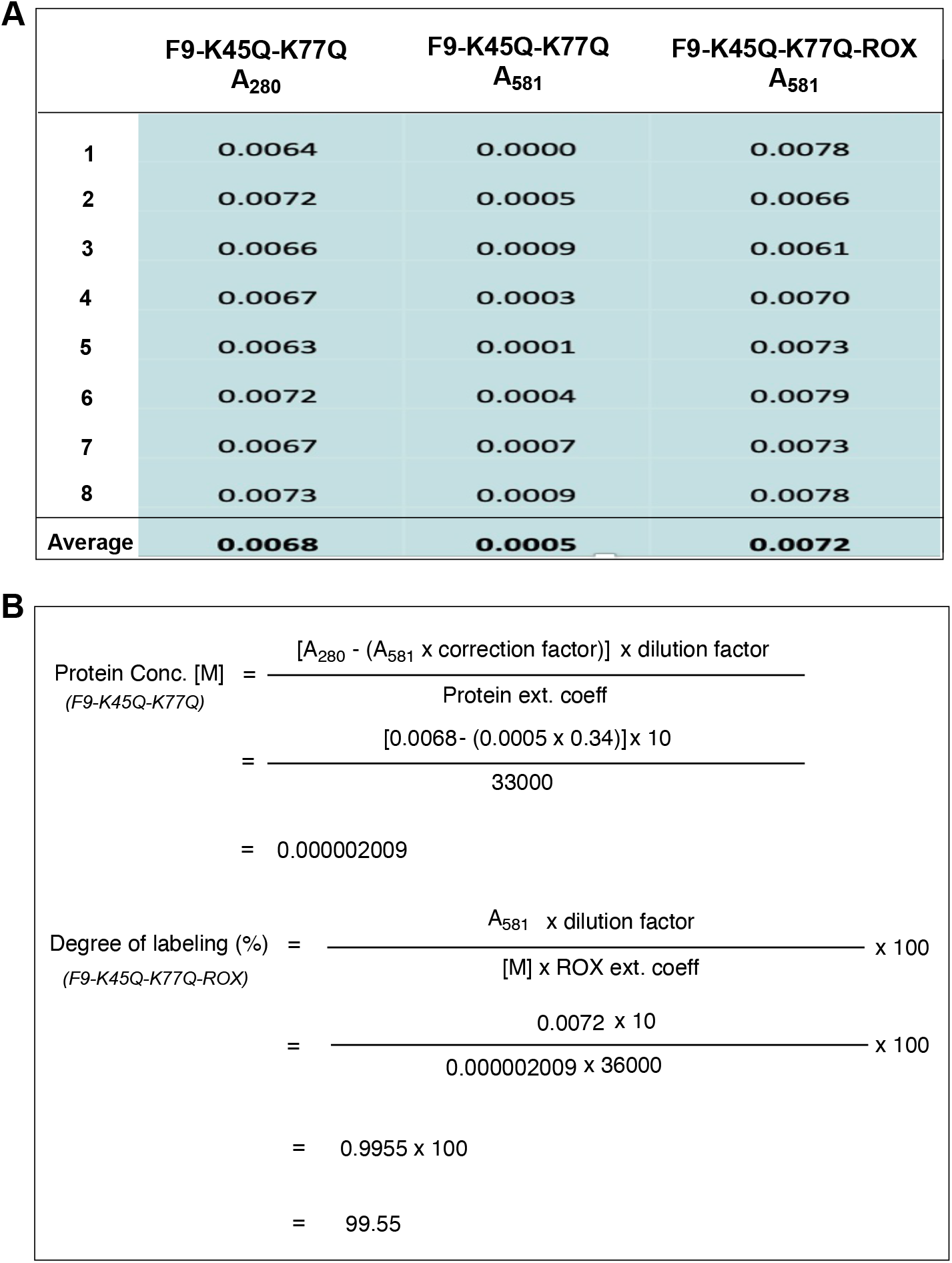
Degree of labeling. (A) Optical density of F9-K45Q-K77Q or F9-K45Q-K77Q-ROX measured by a NanoDrop 3300 Fluorospectrometer at 280 nm and 581 nm. (B) Calculation of degree of labeling from measured by protein concentration.

**Table S1.**
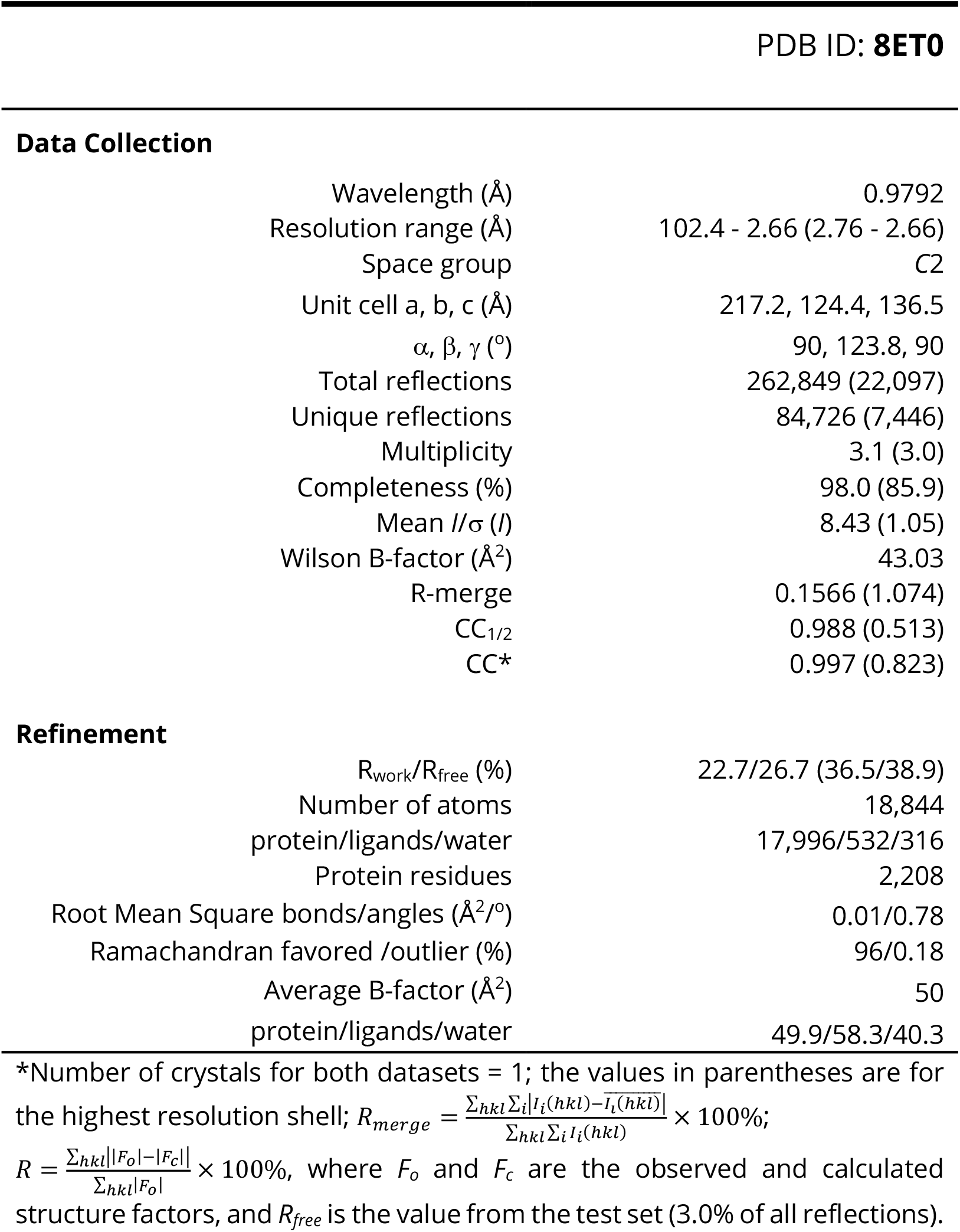
Atomic coordinates and structure factors of mCOX-2:F9 Co-crystal Complex.

## References

1. M. Bacou et al., Development of Preclinical Ultrasound Imaging Techniques to Identify and Image Sentinel Lymph Nodes in a Cancerous Animal Model. Cancers (Basel) 14 (2022).

2. C. Strelez et al., Human colorectal cancer-on-chip model to study the microenvironmental influence on early metastatic spread. iScience 24, 102509 (2021).

3. C. M. Schurch et al., Coordinated Cellular Neighborhoods Orchestrate Antitumoral Immunity at the Colorectal Cancer Invasive Front. Cell 183, 838 (2020).

4. W. M. Grady, C. C. Pritchard, Molecular alterations and biomarkers in colorectal cancer. Toxicol Pathol 42, 124–139 (2014).

5. W. M. Grady, S. D. Markowitz, The molecular pathogenesis of colorectal cancer and its potential application to colorectal cancer screening. Dig Dis Sci 60, 762–772 (2015).

6. E. Koncina, S. Haan, S. Rauh, E. Letellier, Prognostic and Predictive Molecular Biomarkers for Colorectal Cancer: Updates and Challenges. Cancers (Basel) 12 (2020).

7. O. Mazouji, A. Ouhajjou, R. Incitti, H. Mansour, Updates on Clinical Use of Liquid Biopsy in Colorectal Cancer Screening, Diagnosis, Follow-Up, and Treatment Guidance. Front Cell Dev Biol 9, 660924 (2021).

8. A. R. Sepulveda et al., Molecular Biomarkers for the Evaluation of Colorectal Cancer. Am J Clin Pathol 147, 221–260 (2017).

9. M. Goetz, T. D. Wang, Molecular imaging in gastrointestinal endoscopy. Gastroenterology 138, 828–833 e821 (2010).

10. J. Kim et al., Molecular Imaging of Colorectal Tumors by Targeting Colon Cancer Secreted Protein-2 (CCSP-2). Neoplasia 19, 805–816 (2017).

11. B. Bressler et al., Rates of new or missed colorectal cancers after colonoscopy and their risk factors: a population-based analysis. Gastroenterology 132, 96–102 (2007).

12. S. Lin, Y. Li, A. A. Zamyatnin, Jr., J. Werner, A. V. Bazhin, Reactive oxygen species and colorectal cancer. J Cell Physiol 233, 5119–5132 (2018).

13. F. A. Sinicrope, S. Gill, Role of cyclooxygenase-2 in colorectal cancer. Cancer Metastasis Rev 23, 63–75 (2004).

14. K. T. Ng, A. K. V. Tsia, V. Y. L. Chong, Robotic Versus Conventional Laparoscopic Surgery for Colorectal Cancer: A Systematic Review and Meta-Analysis with Trial Sequential Analysis. World J Surg 10.1007/s00268-018-04896-7 (2019).

15. A. T. Abegunde, Chromoendoscopy, White-Light, or Narrow-Band Imaging Colonoscopy in Colorectal Cancer Surveillance in Inflammatory Bowel Disease: True Illumination or Game of Shadows. Clin Gastroenterol Hepatol 14, 1062 (2016).

16. V. Gomez, R. G. Racho, M. G. Heckman, N. N. Diehl, M. B. Wallace, High-definition white-light (HDWL) colonoscopy and higher adenoma detection rate and the potential for paradoxical over surveillance. Dig Dis Sci 59, 2749–2756 (2014).

17. C. J. Kahi et al., High-definition chromocolonoscopy vs. high-definition white light colonoscopy for average-risk colorectal cancer screening. Am J Gastroenterol 105, 1301–1307 (2010).

18. N. S. S. Atkinson et al., Narrow-Band Imaging for Detection of Neoplasia at Colonoscopy: A Meta-analysis of Data From Individual Patients in Randomized Controlled Trials. Gastroenterology 157, 462–471 (2019).

19. S. Zhao et al., Magnitude, Risk Factors, and Factors Associated With Adenoma Miss Rate of Tandem Colonoscopy: A Systematic Review and Meta-analysis. Gastroenterology 10.1053/j.gastro.2019.01.260 (2019).

20. J. C. van Rijn et al., Polyp miss rate determined by tandem colonoscopy: a systematic review. Am J Gastroenterol 101, 343–350 (2006).

21. F. A. Orlando et al., Aberrant crypt foci as precursors in colorectal cancer progression. J Surg Oncol 98, 207–213 (2008).

22. S. Katsuki et al., [Aberrant crypt foci as biomarkers in chemoprevention for colorectal cancer]. Nihon Geka Gakkai Zasshi 99, 379–384 (1998).

23. T. Takayama et al., Aberrant crypt foci of the colon as precursors of adenoma and cancer. N Engl J Med 339, 1277–1284 (1998).

24. N. H. Kim et al., Miss rate of colorectal neoplastic polyps and risk factors for missed polyps in consecutive colonoscopies. Intest Res 15, 411–418 (2017).

25. M. M. Taketo, COX-2 and colon cancer. Inflamm. Res. 47 Suppl 2, S112–116 (1998).

26. L. Moreira, A. Castells, Cyclooxygenase as a target for colorectal cancer chemoprevention. Curr Drug Targets 12, 1888–1894 (2011).

27. S. Samoha, N. Arber, Cyclooxygenase-2 inhibition prevents colorectal cancer: from the bench to the bed side. Oncology 69 Suppl 1, 33–37 (2005).

28. C. H. Koehne, R. N. Dubois, COX-2 inhibition and colorectal cancer. Semin Oncol 31, 12–21 (2004).

29. L. J. Marnett, R. N. DuBois, COX-2: a target for colon cancer prevention. Annu Rev Pharmacol Toxicol 42, 55–80 (2002).

30. M. J. Uddin et al., Selective visualization of cyclooxygenase-2 in inflammation and cancer by targeted fluorescent imaging agents. Cancer Res 70, 3618–3627 (2010).

31. M. J. Uddin, B. C. Crews, K. Ghebreselasie, L. J. Marnett, Design, synthesis, and structure-activity relationship studies of fluorescent inhibitors of cycloxygenase-2 as targeted optical imaging agents. Bioconjugate chemistry 24, 712–723 (2013).

32. M. J. Uddin et al., Trifluoromethyl fluorocoxib a detects cyclooxygenase-2 expression in inflammatory tissues and human tumor xenografts. ACS Med Chem Lett 5, 446–450 (2014).

33. M. J. Uddin et al., Fluorocoxib A loaded nanoparticles enable targeted visualization of cyclooxygenase-2 in inflammation and cancer. Biomaterials 92, 71–80 (2016).

34. M. J. Uddin et al., Molecular Imaging of Inflammation in Osteoarthritis Using a Water-Soluble Fluorocoxib. ACS Med Chem Lett 11, 1875–1880 (2020).

35. M. J. Uddin et al., Fluorocoxib A enables targeted detection of cyclooxygenase-2 in laser-induced choroidal neovascularization. J Biomed Opt 21, 90503 (2016).

36. O. Tietz et al., PET imaging of cyclooxygenase-2 (COX-2) in a pre-clinical colorectal cancer model. EJNMMI Res 6, 37 (2016).

37. C. Tondera et al., Optical imaging of COX-2: studies on an autofluorescent 2,3-diaryl-substituted indole-based cyclooxygenase-2 inhibitor. Biochem Biophys Res Commun 458, 40–45 (2015).

38. H. Zhang et al., An off-on COX-2-specific fluorescent probe: targeting the Golgi apparatus of cancer cells. J Am Chem Soc 135, 11663–11669 (2013).

39. C. Dagallier et al., Development of PET Radioligands Targeting COX-2 for Colorectal Cancer Staging, a Review of in vitro and Preclinical Imaging Studies. Front Med (Lausanne) 8, 675209 (2021).

40. H. M. Schuller et al., Detection of overexpressed COX-2 in precancerous lesions of hamster pancreas and lungs by molecular imaging: implications for early diagnosis and prevention. ChemMedChem 1, 603–610 (2006).

41. C. Hamers-Casterman et al., Naturally occurring antibodies devoid of light chains. Nature 363, 446–448 (1993).

42. H. D. Herce et al., Cell-permeable nanobodies for targeted immunolabelling and antigen manipulation in living cells. Nat Chem 9, 762–771 (2017).

43. M. A. de Beer, B. N. G. Giepmans, Nanobody-Based Probes for Subcellular Protein Identification and Visualization. Front Cell Neurosci 14, 573278 (2020).

44. P. Bannas, J. Hambach, F. Koch-Nolte, Nanobodies and Nanobody-Based Human Heavy Chain Antibodies As Antitumor Therapeutics. Front Immunol 8, 1603 (2017).

45. T. J. Harmand, A. Islam, N. Pishesha, H. L. Ploegh, Nanobodies as in vivo, non-invasive, imaging agents. RSC Chem Biol 2, 685–701 (2021).

46. I. Van Audenhove, J. Gettemans, Nanobodies as Versatile Tools to Understand, Diagnose, Visualize and Treat Cancer. EBioMedicine 8, 40–48 (2016).

47. D. R. Maass, J. Sepulveda, A. Pernthaner, C. B. Shoemaker, Alpaca (Lama pacos) as a convenient source of recombinant camelid heavy chain antibodies (VHHs). J Immunol Methods 324, 13–25 (2007).

48. C. Vincke et al., General strategy to humanize a camelid single-domain antibody and identification of a universal humanized nanobody scaffold. J Biol Chem 284, 3273–3284 (2009).

49. S. W. Rowlinson et al., A novel mechanism of cyclooxygenase-2 inhibition involving interactions with Ser-530 and Tyr-385. J. Biol. Chem. 278, 45763–45769 (2003).

50. M. J. Uddin et al., Antitumor Activity of Cytotoxic Cyclooxygenase-2 Inhibitors. ACS Chem Biol 11, 3052–3060 (2016).

51. T. Bartoschik et al., Near-native, site-specific and purification-free protein labeling for quantitative protein interaction analysis by MicroScale Thermophoresis. Sci Rep 8, 4977 (2018).

52. K. C. Duggan et al., Molecular basis for cyclooxygenase inhibition by the non-steroidal anti-inflammatory drug naproxen. J. Biol. Chem. 285, 34950–34959 (2010).

53. D. Dzhalilova, N. Zolotova, N. Fokichev, O. Makarova, Murine models of colorectal cancer: the azoxymethane (AOM)/dextran sulfate sodium (DSS) model of colitis-associated cancer. PeerJ 11, e16159 (2023).

54. Q. Pan et al., Genomic variants in mouse model induced by azoxymethane and dextran sodium sulfate improperly mimic human colorectal cancer. Sci Rep 7, 25 (2017).

55. R. Suzuki, H. Kohno, S. Sugie, T. Tanaka, Sequential observations on the occurrence of preneoplastic and neoplastic lesions in mouse colon treated with azoxymethane and dextran sodium sulfate. Cancer Sci 95, 721–727 (2004).

56. D. Kalyane et al., Employment of enhanced permeability and retention effect (EPR): Nanoparticle-based precision tools for targeting of therapeutic and diagnostic agent in cancer. Mater Sci Eng C Mater Biol Appl 98, 1252–1276 (2019).

57. M. J. Uddin et al., Selective visualization of cyclooxygenase-2 in inflammation and cancer by targeted fluorescent imaging agents. Cancer Res. 70, 3618–3627 (2010).

58. M. J. Uddin et al., Targeted imaging of cancer by fluorocoxib C, a near-infrared cyclooxygenase-2 probe. J Biomed Opt 20, 50502 (2015).

59. M. J. Uddin, B. C. Crews, K. Ghebreselasie, L. J. Marnett, Design, synthesis, and structure-activity relationship studies of fluorescent inhibitors of cycloxygenase-2 as targeted optical imaging agents. Bioconjug. Chem. 24, 712–723 (2013).

60. Y. C. Huang, Y. C. Chang, C. N. Yeh, C. S. Yu, Synthesis and Biological Evaluation of an (18)Fluorine-Labeled COX Inhibitor--[(18)F]Fluorooctyl Fenbufen Amide--For Imaging of Brain Tumors. Molecules 21, 387 (2016).

61. T. Toyokuni et al., Synthesis of 4-(5-[18F]fluoromethyl-3-phenylisoxazol-4-yl)benzenesulfonamide, a new [18F]fluorinated analogue of valdecoxib, as a potential radiotracer for imaging cyclooxygenase-2 with positron emission tomography. Bioorg Med Chem Lett 15, 4699–4702 (2005).

62. M. J. Uddin, H. Niitsu, R. J. Coffey, L. J. Marnett, Development of Pluoronic nanoparticles of fluorocoxib A for endoscopic fluorescence imaging of colonic adenomas. J Biomed Opt 28, 040501 (2023).

63. E. Pardon et al., A general protocol for the generation of Nanobodies for structural biology. Nat Protoc 9, 674–693 (2014).

64. S. W. Rowlinson et al., A novel mechanism of cyclooxygenase-2 inhibition involving interactions with Ser-530 and Tyr-385. J Biol Chem 278, 45763–45769 (2003).

65. S. W. Rowlinson, B. C. Crews, C. A. Lanzo, L. J. Marnett, The binding of arachidonic acid in the cyclooxygenase active site of mouse prostaglandin endoperoxide synthase-2 (COX-2). A putative L-shaped binding conformation utilizing the top channel region. J Biol Chem 274, 23305–23310 (1999).

66. M. M. Mitchener et al., Competition and allostery govern substrate selectivity of cyclooxygenase-2. Proc Natl Acad Sci U S A 112, 12366–12371 (2015).

67. W. Kabsch, Xds. Acta Crystallogr. D Biol. Crystallogr. 66, 125–132 (2010).

68. A. J. McCoy, Solving structures of protein complexes by molecular replacement with Phaser. Acta Crystallogr. D Biol. Crystallogr. 63, 32–41 (2007).

69. P. D. Adams et al., PHENIX: building new software for automated crystallographic structure determination. Acta Crystallogr D Biol Crystallogr 58, 1948–1954 (2002).

